# Proteome-wide serology reveals immune-defined subtypes of gastrointestinal disease in systemic sclerosis

**DOI:** 10.64898/2026.05.19.724137

**Authors:** Zsuzsanna H. McMahan, Srinivas Puttapaka, Tyler Hulett, Ami A. Shah, Kathryn Faheem, Shaohui Hu, Gamze Sonmez, Pedro Ramos, Subhash Kulkarni

**Author notes:** **Contact authors:** Zsuzsanna H McMahan, MD, Subhash Kulkarni, PhD.

## Abstract

**Background:** Gastrointestinal (GI) involvement in systemic sclerosis (SSc) affects up to 90% of patients and is a major driver of morbidity and mortality. Despite its clinical importance, GI disease in SSc is highly heterogeneous, with upper and lower GI manifestations representing distinct phenotypic extremes whose underlying immunologic basis remains poorly defined.

**Methods:** We performed unbiased, proteome-wide autoantibody profiling using a human protein microarray comprising >21,000 full-length proteins (>80% of the human proteome). Sera from patients with SSc and isolated upper GI dysmotility (n=23), isolated lower GI dysmotility (n=17), and non-SSc controls (n=20) were analyzed. Enriched autoantibodies were identified using Fisher’s exact test, and unsupervised clustering was applied to define serology-based patient subsets and relate immune signatures to clinical phenotypes.

**Results:** Distinct autoantibody profiles differentiated patients with upper versus lower GI disease. Upper GI–predominant SSc was characterized by enrichment of previously unreported autoantibodies, including those targeting TiSSc1/2 (newly identified proteins encoded within the MIRLET7BHG locus), FAM9C, SPATA20, FAM110D, EMILIN1, CARD14, SMN1, KCTD7, and PHYHD1, whereas lower GI disease was associated with antibodies against HAO2, KLHL7, SUFU, APPL1, BNIP2, UCHL3, ZNF385A, LIMD1, MAGEA9, and PPP2R3C. Serology-driven clustering identified four reproducible subgroups with distinct patterns of GI, pulmonary, vascular, and autonomic involvement, defining clinically meaningful disease phenotypes that extend beyond traditional anatomic classification.

**Conclusions:** Proteome-scale serological profiling reveals previously unrecognized autoimmune signatures underlying GI heterogeneity in SSc. These findings support a shift from anatomy-based to serology-defined classification of SSc GI disease and provide a foundation for biomarker development, patient stratification, and precision medicine approaches in this population.

## INTRODUCTION

Systemic sclerosis (SSc) is a complex autoimmune disease characterized by fibrosis, vasculopathy, and immune dysregulation affecting both cutaneous tissues and visceral organs^111^. Among these, the gastrointestinal (GI) tract is particularly impacted, with dysfunction affecting up to 90% of SSc patients^2^. The severity of GI involvement, ranging from esophageal to colonic dysmotility, is associated with increased mortality^2,3^. Complications including malnutrition^4,5^, intestinal failure^6,7^, and refractory symptoms due to dysmotility^8^ substantially impair quality of life and clinical outcomes^9^, underscoring the critical importance of GI disease in determining SSc trajectory.

GI manifestations in SSc are highly heterogenous and evolve over time^10^. Although prior studies have identified clinical features and serological factors associated with GI severity^10–12^, regional involvement, and disease progression^10^, the biological drivers of this variability remain poorly understood. A central hypothesis is that SSc comprises biologically distinct subtypes shaped by discrete pathogenic processes. In this framework, patients with predominantly upper GI involvement (esophageal and/or gastric dysfunction) versus those with predominantly lower GI involvement (small intestinal and/or colonic dysfunction) represent two clinically divergent phenotypes that may arise from region-specific mechanisms.

These differences may reflect variation in underlying neural regulation^13^ and tissue biology^14^. Upper GI motility is thought to be regulated by actions of motor vagal neurons in concert with the enteric nervous system (ENS), whereas the contribution of vagal neurons to regulation of small bowel and colonic motility is thought to be minimal and motility in these organs is thought to depend primarily on ENS circuits^13^. The internal anal sphincter is regulated by both ENS and sacral innervation^13^. These anatomically and functionally distinct control systems, together with the regional tissue biology, suggest that autoimmune targeting in SSc may vary by GI segment. Consequently, differences in GI phenotype may correspond to distinct autoantibody profiles and underlying immune programs. Defining phenotype-specific autoantigen repertoires is therefore essential to understanding SSc⍰associated GI dysfunction, yet progress has been limited by the absence of a comprehensive approaches to characterize the autoantigen-ome.

Multiple groups, including our own, have identified novel autoantibodies in SSc^15–17^. Importantly, antibody⍰mediated mechanisms of GI dysfunction are increasingly recognized, with anti-muscarinic receptor type 3 (M3R) and anti⍰mitochondrial M2 antibodies associated with specific clinical phenotypes and patterns of GI involvement^14,18,19^. These findings support the concept that SSc GI disease may be driven by diverse, and potentially region-specific, autoantibody repertoires rather than a single pathogenic specificity. However, conventional approaches for antibody discovery are limited by low throughput and restricted antigen coverage, typically identifying only a small number of targets at a time^15^. Although newer multiplex platforms allow broader profiling, they remain constrained to predefined antigen sets^20,21^. As a result, it remains unclear whether comprehensive, unbiased interrogation of the autoantigen-ome can define immune signatures that better explain GI heterogeneity in SSc.

In this study, we performed proteome-wide autoantigen profiling using a human protein (HuProt) microarray comprising over 21,000 correctly folded proteins^22^, to test whether distinct GI phenotypes in SSc are associated with distinct autoantibody signatures. We analyzed sera from patients with isolated upper or lower GI dysmotility and identified phenotype-specific autoantibody repertoires, including previously unrecognized targets. Integrative analysis further revealed serology-defined patient subsets with distinct clinical and physiological features. Together, these findings provide a systems-level view of the SSc autoantigen-ome and establish a framework for serology-based stratification of GI disease.

## METHODS

### Patient groups

Sera were obtained from the Gastrointestinal Assessment Protocol (GAP) within the Johns Hopkins Scleroderma Center Research Registry, which includes demographic and clinical data collected at baseline and at 6-month intervals, as previously described ^11,14,23,24^. All patients underwent whole gut transit studies assessing motility from the esophagus to the colon. Serum was collected from all patients in the cohort with only upper GI dysmotility (n=23; *upper GI group*) and all patients in the cohort with only lower GI dysmotility (n=17; *lower GI group*) as defined by whole gut scintigraphy. Serum from 20 age-matched healthy controls (*control group*) was also included. All protocols were approved by the Johns Hopkins Institutional Review Board, and participants provided informed consent. Patient-reported GI symptom burden was assessed using the maximum University of California Los Angeles Scleroderma Clinical Trial Consortium GI Tract Instrument 2.0 (UCLA GIT 2.0) survey score recorded during follow-up.

### HuProt Human Proteome Microarray

Serum autoantibody profiling was performed using the HuProt Human Proteome Microarray v4.0 (CDI Laboratories, Mayaguez, PR) as previously described^25,26^. To enable unbiased discovery of novel autoantibodies in SSc-GID, this platform, which comprises >21,000 unique, full-length human proteins and isoforms (covering >81% of the human proteome) was interrogated in duplicate, with 2,000 technical replicates. Net signal intensity for each antigen was defined as the background-subtracted median spot intensity, log₂-transformed, and normalized to the median total signal intensity per sample to account for inter-individual variability in baseline autoantibody levels.

### Analyses of HuProt data

To identify phenotype-enriched autoantibody signatures, we performed parallel pairwise comparisons of upper and lower GI disease groups versus controls, rather than a single multi-group analysis. This approach was selected to maximize sensitivity for detecting phenotype-specific signals that could be obscured in simultaneous multi-group testing, particularly within a high-dimensional feature space and modest-sized cohorts. For each comparison, enriched autoantibody profiles were identified using Fisher’s exact test (P < 0.05) and subsequently subjected to unsupervised clustering to define putative patient subgroups sharing common serological signatures. Clustering analyses were conducted independently using upper GI- and lower GI-enriched feature sets to assess whether distinct analytical frameworks converged on similar subgroup architectures, thereby providing an internal measure of robustness. This strategy additionally enabled independent derivation of phenotype-associated feature sets and evaluation of the reproducibility of clustering patterns across complementary analyses. Disease group-specific autoantibodies were defined as those enriched within a given SSc-GID subgroup and detected in ≤5% of control participants. Collectively, these analyses enabled characterization of subgroup-enriched autoantibody signatures, as well as autoantibodies preferentially detected in control individuals.

### Cell culture experiments

HepG2 (well-differentiated human hepatocellular carcinoma) and LX-2 (immortalized human activated hepatic stellate; kind gift from Prof. Scott Friedman, Icahn School of Medicine at Mt Sinai) cell lines were maintained in Dulbecco’s modified Eagle medium (DMEM; Thermo Fisher Scientific) supplemented with 10% fetal bovine serum (FBS; Sigma-Aldrich), 100 units/mL penicillin/streptomycin, and 4 mM L-glutamine (Thermo Fisher Scientific) at 37°C in 5% CO₂. Media were refreshed every 72 hours. Cells were passaged at approximately 80% confluence using trypsin (Thermo Fisher Scientific) and re-seeded at a 1:5 split ratio.

### Exposure to Heat Shock

LX-2 cells were cultured in complete medium for 24 hours before heat shock. Hyperthermia exposure was performed as previously described^27^, with minor modifications. Briefly, cells were transferred into 1.5 mL tubes, sealed, and incubated in a dry heat block at either 37°C (control) or 43 - 44°C (heat shock) for 20 min. Cells were then replated in 24-well plates and allowed to recover at 37°C in 5% CO₂ for 24 hours prior to downstream analyses.

### Immunofluorescence Staining

Cells were fixed with freshly prepared 1% paraformaldehyde (PFA) for 5 min at room temperature, washed three times with phosphate-buffered saline (PBS), and permeabilized and blocked in PBS containing 5% normal goat serum and 0.5% Triton X-100 for 30 min. Cells were incubated overnight at 16°C with rabbit polyclonal FLJ27365 antibody (Biorbyt, orb477598; 1:500). After washing, cells were incubated with goat anti-rabbit Alexa Fluor 647 secondary antibody (1:500) for 1 hour at room temperature, followed by nuclear staining with DAPI. Coverslips were mounted using DAPI-containing mounting medium. Images were acquired using an EVOS fluorescence microscope (LX-2) or a Leica confocal microscope (HepG2). Mean fluorescence intensity (MFI) was quantified in ImageJ by defining the total cellular region of interest for each cell. Analyses included at least three independent experiments with at least 30 cells quantified per replicate

### RNA isolation and Quantitative real time PCR

Total RNA was extracted from LX2 cells following exposure to physiological temperature or hyperthermia (as detailed above) using Direct-zol RNA Miniprep Kit (Zymo research). One microgram of RNA was reverse transcribed into cDNA using iScript™ Reverse Transcription Supermix (Biorad) in a 20μl reaction. Quantitative real-time PCR was performed on a QuantStudio3 real time PCR system (Life Technologies, USA) using Powerstrack SYBR Green Master Mix (Applied Biosystems). Primer sequences are in **Table 1**. Relative gene expression was calculated using the 2^−ΔΔCt^ method.

**Table 1:**
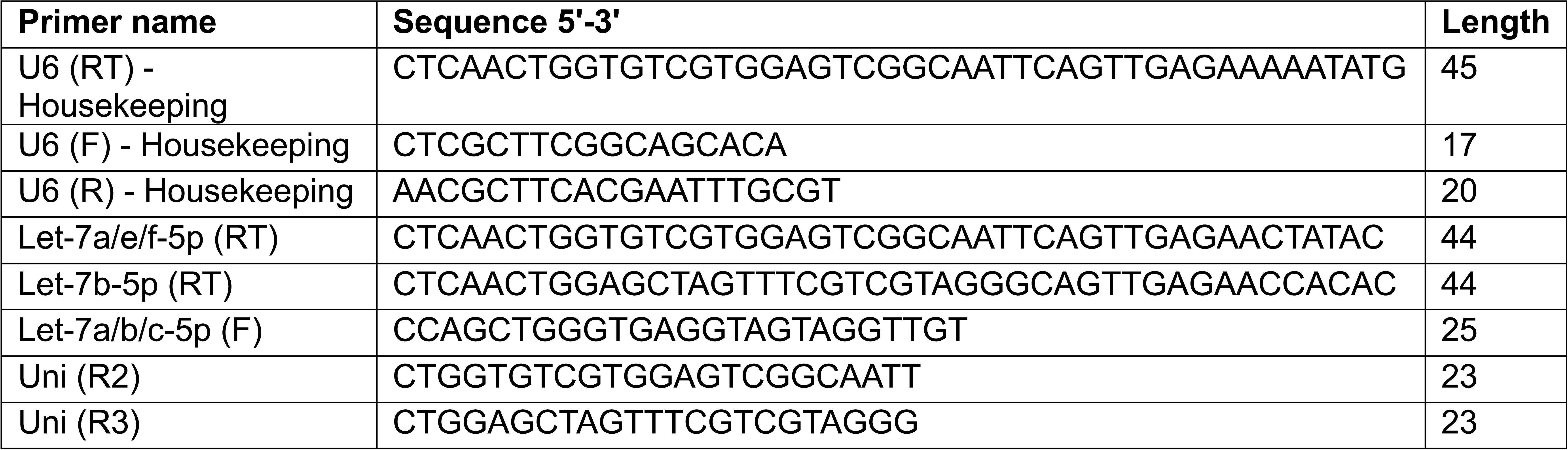
Primer names and sequences used for quantitative real time PCR.

## RESULTS

### Distinct serological signatures define upper GI involvement in SSc

We compared normalized HuProt signals from patients with upper GI involvement to those from all remaining participants (lower GI disease and controls). Using Fisher’s exact test (p ≤ 0.05), we identified a set of autoantibodies significantly enriched in upper GI disease. These are visualized in **Fig. 1**, where signal intensity reflects relative antibody titers in patients with upper GI disease (n = 23) compared with the remainder of the cohort (n = 37). The prevalence of each autoantibody across groups is summarized in **Table 2**.

**Fig. 1:**
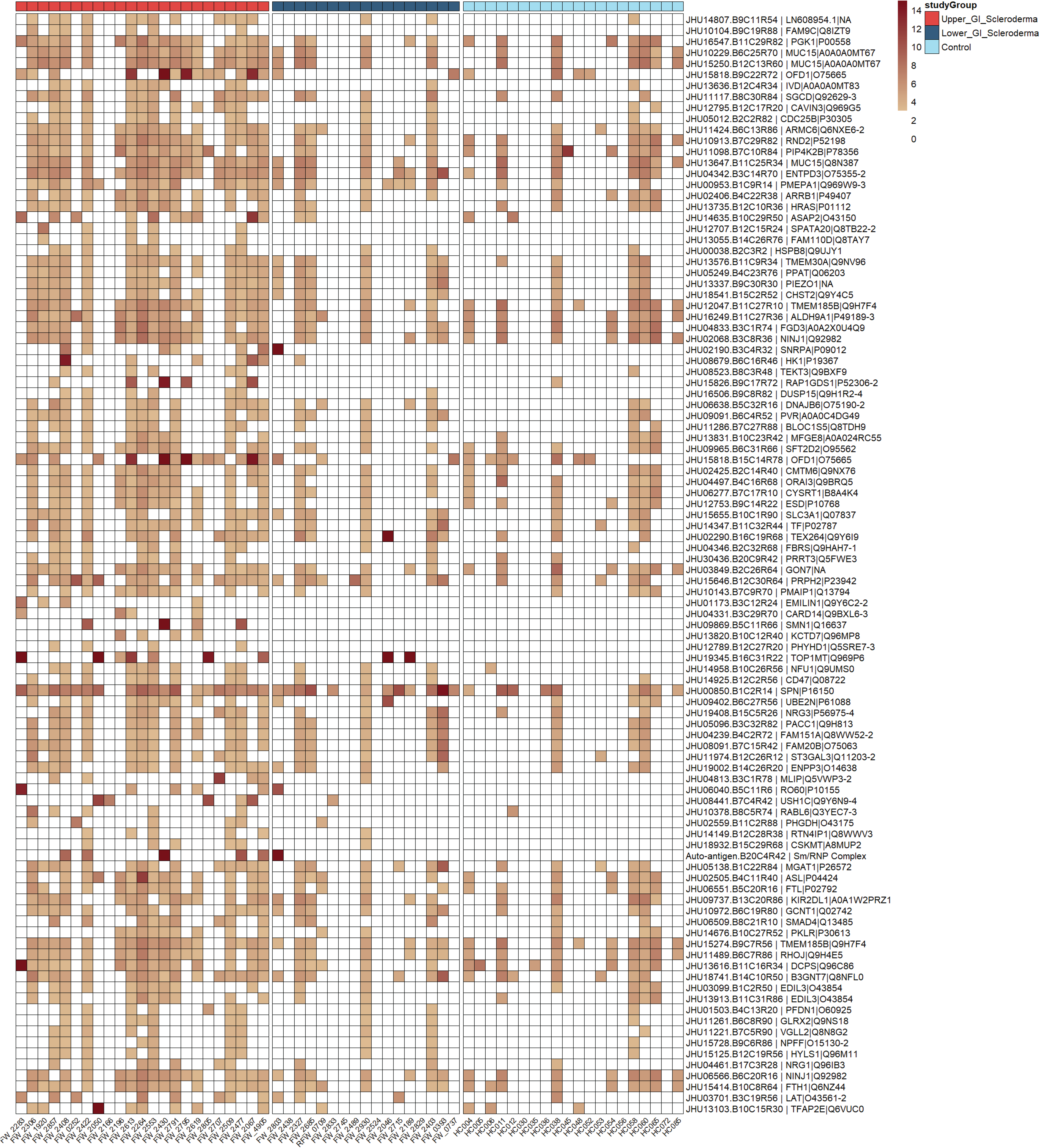
Autoantigens enriched in SSc patients with upper GI disease. Heatmap showing proteins targeted by autoantibodies significantly enriched in patients with SSc and upper GI involvement (red) compared with the remainder of the cohort, including patients with lower GI disease (dark blue) and non-SSc controls (light blue). Columns represent individual participants (patient IDs on the x-axis), and rows represent autoantigens. Color intensity reflects relative autoantibody signal. Only autoantibodies meeting statistical significance (Fisher’s exact test, pL<L0.05; upper GI vs all other participants) are shown.

**Table 2:**
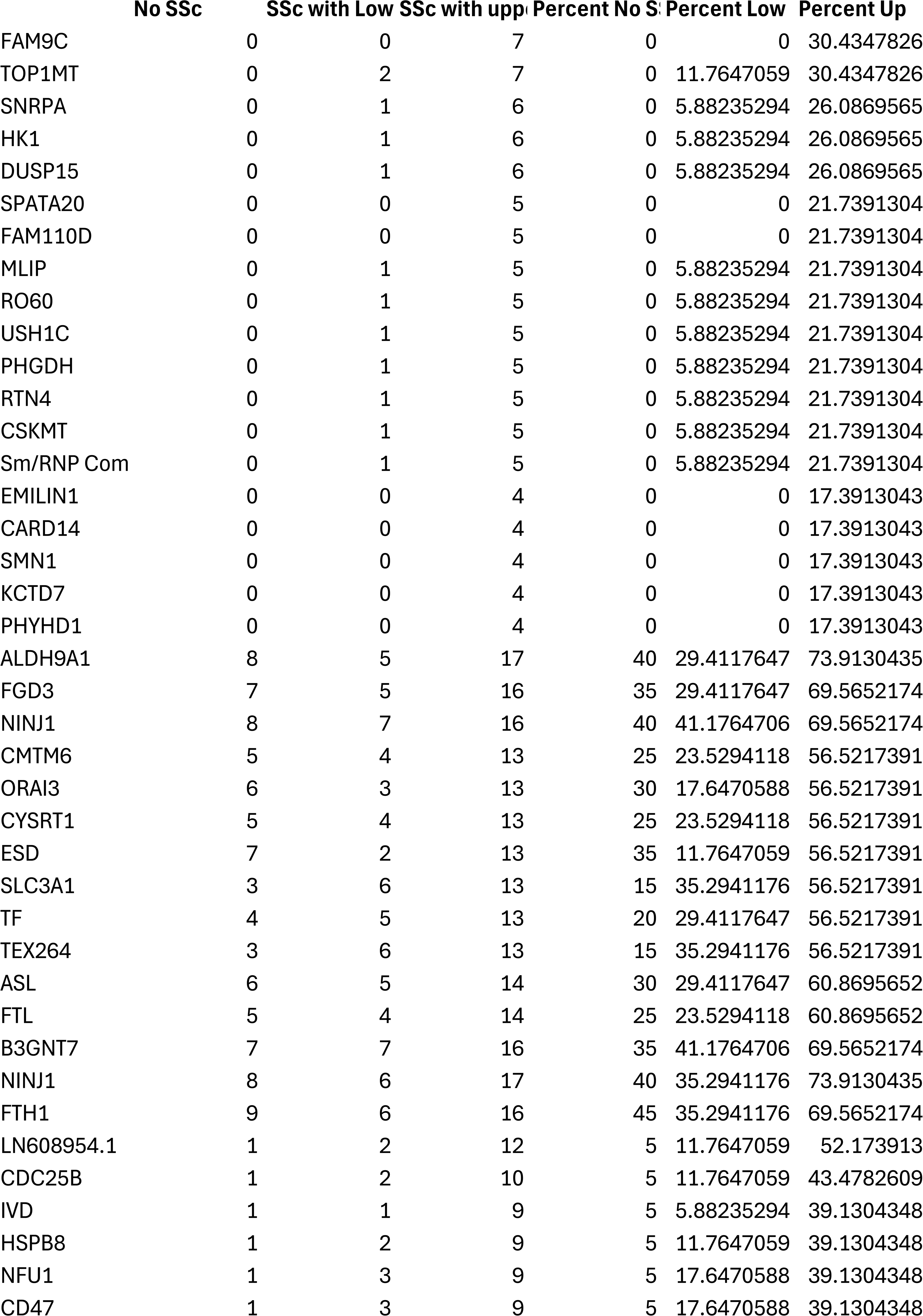
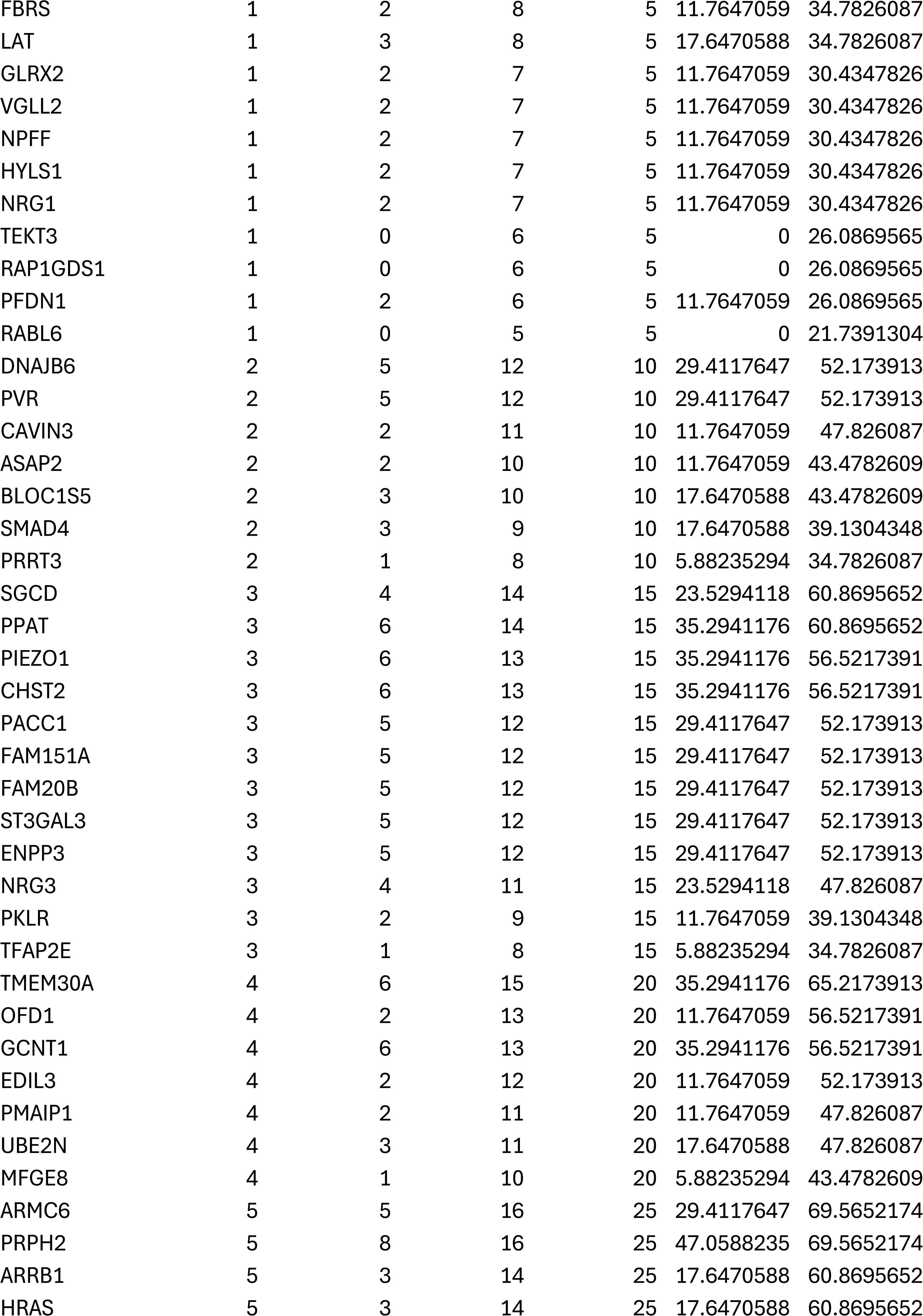
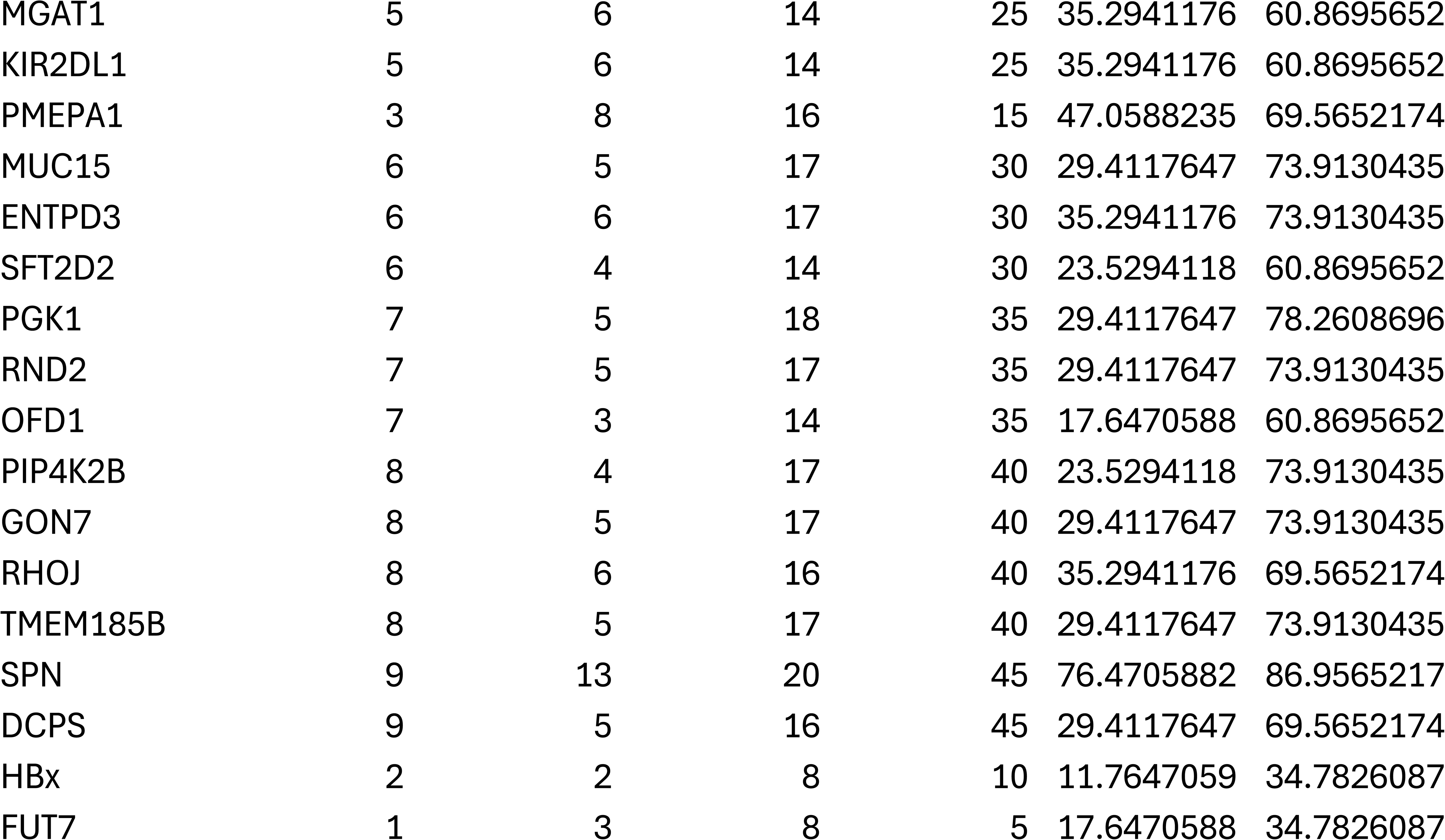
Proportion of patients with upper GI–predominant SSc, lower GI–predominant SSc, and non-SSc controls exhibiting autoantibodies against antigens significantly enriched in upper GI–associated SSc compared with the other cohorts.

This analysis identified multiple autoantibodies uniquely detected in upper GI disease and absent in both lower GI disease and controls, including those targeting FAM9C (30.43%), SPATA20 (21.74%), FAM110D (21.74%), EMILIN1, CARD14, SMN1, KCTD7, and PHYHD1 (each 17.39%). To our knowledge, these specificities have not previously been described in SSc. Additional autoantibodies (including those targeting SNRPA, HK1, and DUSP15, present in 26.08% of upper GI patients, and MLIP, RO60, USH1C, PHGDH, RTN, CSKMT, and the Sm/RNP complex, present in 21.74%) were absent in controls but detected at low frequency (5.88%) in patients with lower GI disease, indicating SSc-specific enrichment with marked predominance in upper GI phenotypes.

Upper GI disease was further characterized by high-prevalence enrichment of autoantibodies targeting LN608954.1 (52.17%) and CDC25B (43.47%), followed by IVD, HSPB8, NFU, and CD47 (39.13%), and FUT7 (34.78%). These signals were substantially less frequent in lower GI disease (<20% in most cases), reinforcing phenotype-specific enrichment. Notably, antibodies against hepatitis B virus protein X (HBx) were more prevalent in upper GI disease (34.78%) compared with lower GI disease (11.76%) and controls (10%).

### Identification and characterization of a novel autoantigen (TiSSc)

Approximately half of patients with upper GI disease exhibited autoantibodies against LN608954.1, a previously unannotated protein. As shown **in Fig. 2A(i)**, LN608954.1 encodes a 160–amino acid protein, which we designated Targeted in Systemic Sclerosis 1 (TiSSc1). Genomic analyses demonstrated that TiSSc1 is encoded within the MIRLET7BHG locus (22q13.31), which also hosts the *let*⍰l*7* microRNAs (*let*⍰l*7a* and *let*⍰l*7b*)^28^ (**Supplementary Fig. 1A**). An alternatively spliced variant encoding a distinct 219–amino acid protein (TiSSc2) was identified (**Fig. 2A(ii)**), sharing the N-terminal sequence with TiSSc1 but diverging thereafter (**Supplementary Fig. 1A**). Genomic mapping revealed that TiSSc2-specific exons are embedded within intronic regions of the TiSSc1 locus, with *let*⍰l*7* microRNAs located in the 3′ untranslated regions (**Supplementary Fig. 1B**).

**Fig. 2:**
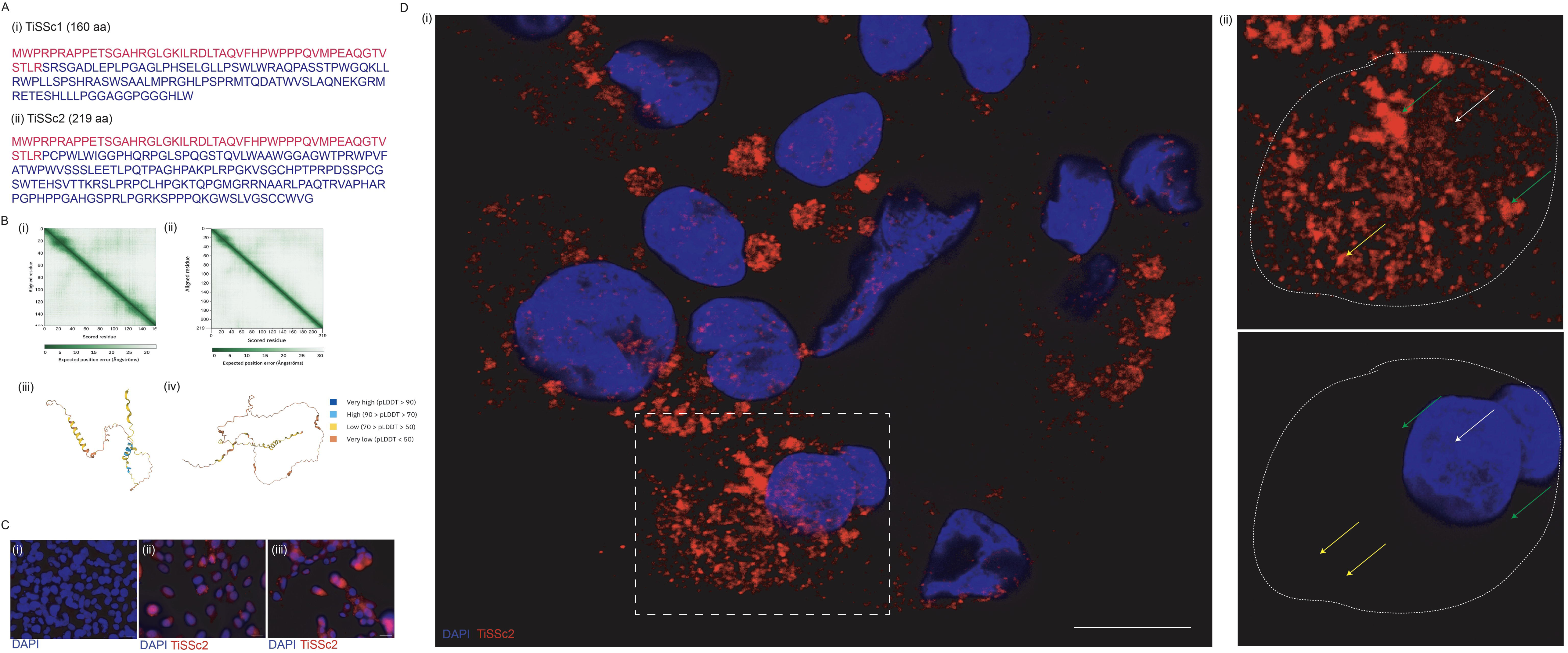
Characterization of the novel autoantigen Targeted in Systemic Sclerosis (TiSSc1/2). (A) Amino acid sequences of (i) TiSSc1 (160 aa) and (ii) TiSSc2 (219 aa), with conserved regions shown in red and divergent regions in blue. (B) AlphaFold structural predictions for TiSSc1 (i, iii) and TiSSc2 (ii, iv). Predicted aligned error (PAE) indicates consistent positional relationships, whereas low predicted local distance difference test (pLDDT) scores (<70) across most residues indicate low confidence in ordered structure. (C) Immunofluorescence staining of HepG2 and LX-2 cells using a TiSSc2-specific antibody (red). (i–ii) Representative images show robust intracellular staining; (iii) unstained LX-2 cells confirm specificity. Nuclei are counterstained with DAPI (blue). Images were acquired with a 40× objective (EVOS microscope). Scale bar, 10Lµm. (D) (i) Confocal imaging of HepG2 cells stained for TiSSc2 (red) and DAPI (blue) (63× oil immersion). Scale bar, 10Lµm. (ii) Magnified view of the boxed region in (i), showing cytoplasmic enrichment with focal nuclear signal (white arrow), perinuclear clustering (green arrow), and diffuse cytoplasmic distribution (yellow arrows).

Comparative genomic analyses and codonization indicated that TiSSc represents an evolutionarily recent gene with conservation restricted to Old World monkeys^29,30^ (**Supplementary Figs. 2–4**). Structural predictions of both TiSSc1 and TiSSc2 using AlphaFold^31,32^ (**Fig. 2B**) showed very low confidence when predicting the protein’s ordered structure, given that the Predicted Local Distance Difference Test (pLDDT) values for most of the structure fell below 70.

Publicly available mass spectrometry datasets (OpenProt^33^) provided independent evidence of the presence of TiSSc1-derived peptides in human tissues (**Supplementary Fig. 5**). We experimentally verified the presence of TiSSc2 by performing immunocytochemistry using a TiSSc2-specific (against 139-219 residues) antibody and observed predominant cytoplasmic localization in LX⍰2 and HepG2 cells (**Fig. 2C**), with additional nuclear signal detected by confocal microscopy (**Fig. 2D**). We then tested whether cellular stress altered the abundance of TiSSc2. Exposure to hyperthermia (43–44°C) significantly increased TiSSc2 abundance (**Supplementary Fig. 6A;** p = 0.0075), whereas *let*⍰l*7a/b* expression remained unchanged (**Supplementary Fig. 6 B, C**), suggesting stress-responsive protein regulation independent of MIRLET7BHG transcription.

Given shared N-terminal homology, we conservatively classified autoantibodies against LN608954.1 as anti⍰TiSSc1/2. Individuals harboring these autoantibodies demonstrated delayed esophageal transit (**Supplementary Table 1;** p = 0.01) consistent with their strong association with upper GI disease. These patients also showed a trend toward male sex, elevated Right Ventricular Systolic Pressure (RVSP), and co-positivity for anti-centromere autoantibodies, although these findings did not reach statistical significance.

### Distinct autoantibody profiles define lower GI disease in SSc

We next compared normalized HuProt signals from patients with lower GI disease to those from the rest of the patients (upper GI disease and controls). Using Fisher’s exact test (p ≤ 0.05), we identified a distinct set of autoantibodies enriched in lower GI disease (**Fig. 3**; **Table 3**). Several autoantibodies were uniquely detected in lower GI disease, including those targeting HAO2, KLHL7, SUFU, and APPL1 (each 23.53%), as well as BNIP2, UCHL3, ZNF385A, LIMD1, MAGEA9, CARHSP1, PPP2R3C, and GPCPD1 (each 17.64%). Consistent with prior observations, antibodies against AGO2 were also enriched in lower GI disease^36^. Additional enriched targets included NFKBIB (35.29%) and SUGT1 (29.41%), along with RTKN2, POP1, MACROH2A2, STX16, and RPP25 (each 23.53%), all of which were absent in controls and in less than 10% of upper GI disease patients. In addition, we identified several novel autoantibodies enriched specifically in patients with lower GI disease and present in less than 5% of patients in both the upper GI disease and control groups. These included autoantibodies directed against NSL1 and FYTTD1 (each in 29.41% patients), as well as CTNNA2, FOLH1, and NEDD4L (each in 23.53% of patients). Autoantibodies against centromere proteins (CENPA, CENPB, CENPC, CENPP) were absent in controls but more frequently detected in lower GI disease than in upper GI disease patients (**Table 3**), consistent with prior associations between centromere serology and GI involvement^34–38^.

**Fig. 3:**
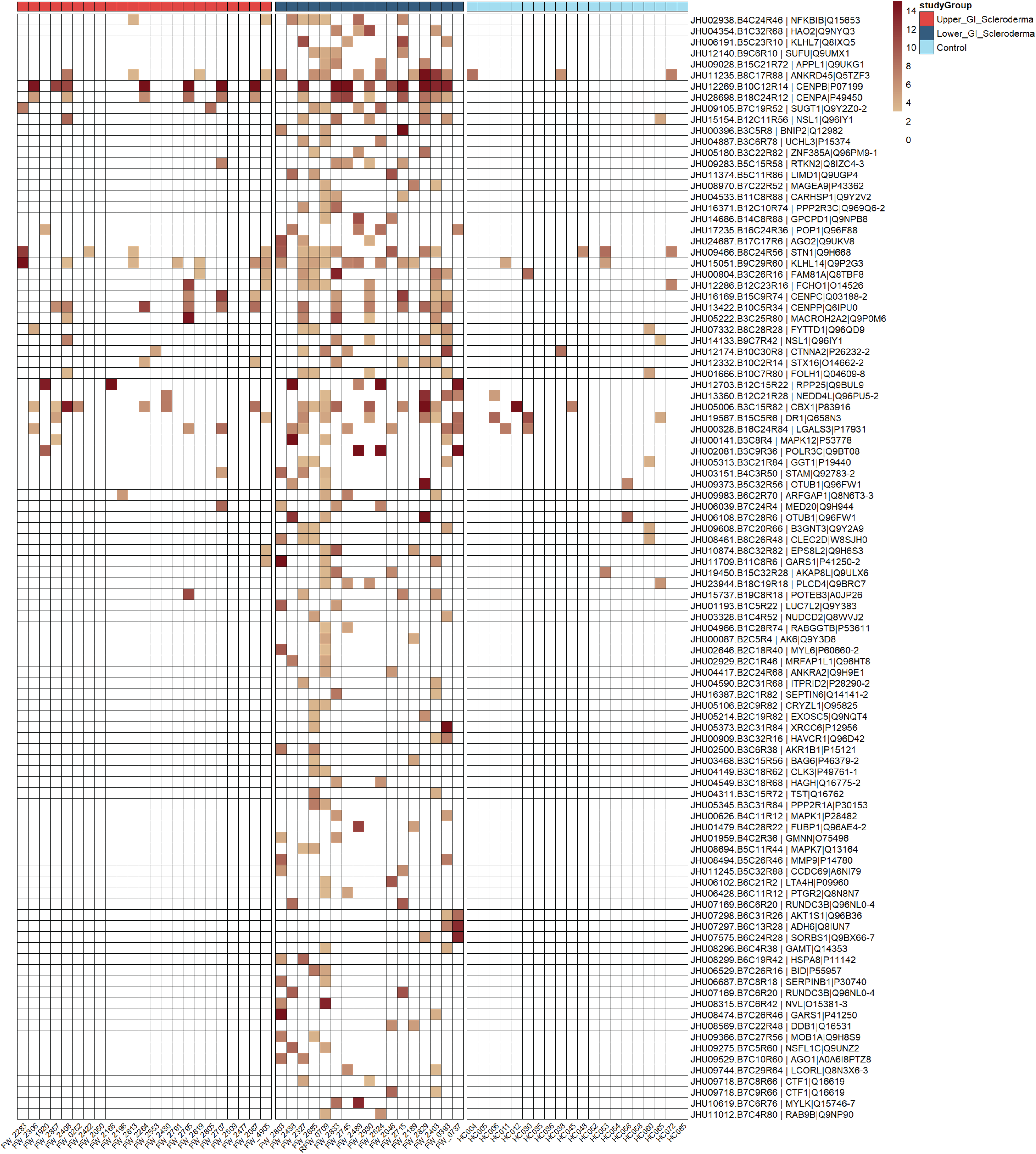
Autoantibodies enriched in SSc patients with lower GI disease. Heatmap showing proteins targeted by autoantibodies significantly enriched in patients with SSc and lower GI involvement (dark blue) relative to patients with upper GI disease (red) and controls (light blue). Columns represent individual participants and rows represent autoantigens. Color intensity reflects relative autoantibody signal. Only statistically significant autoantibodies (Fisher’s exact test, pL<L0.05; lower GI vs all others) are displayed.

**Table 3:**
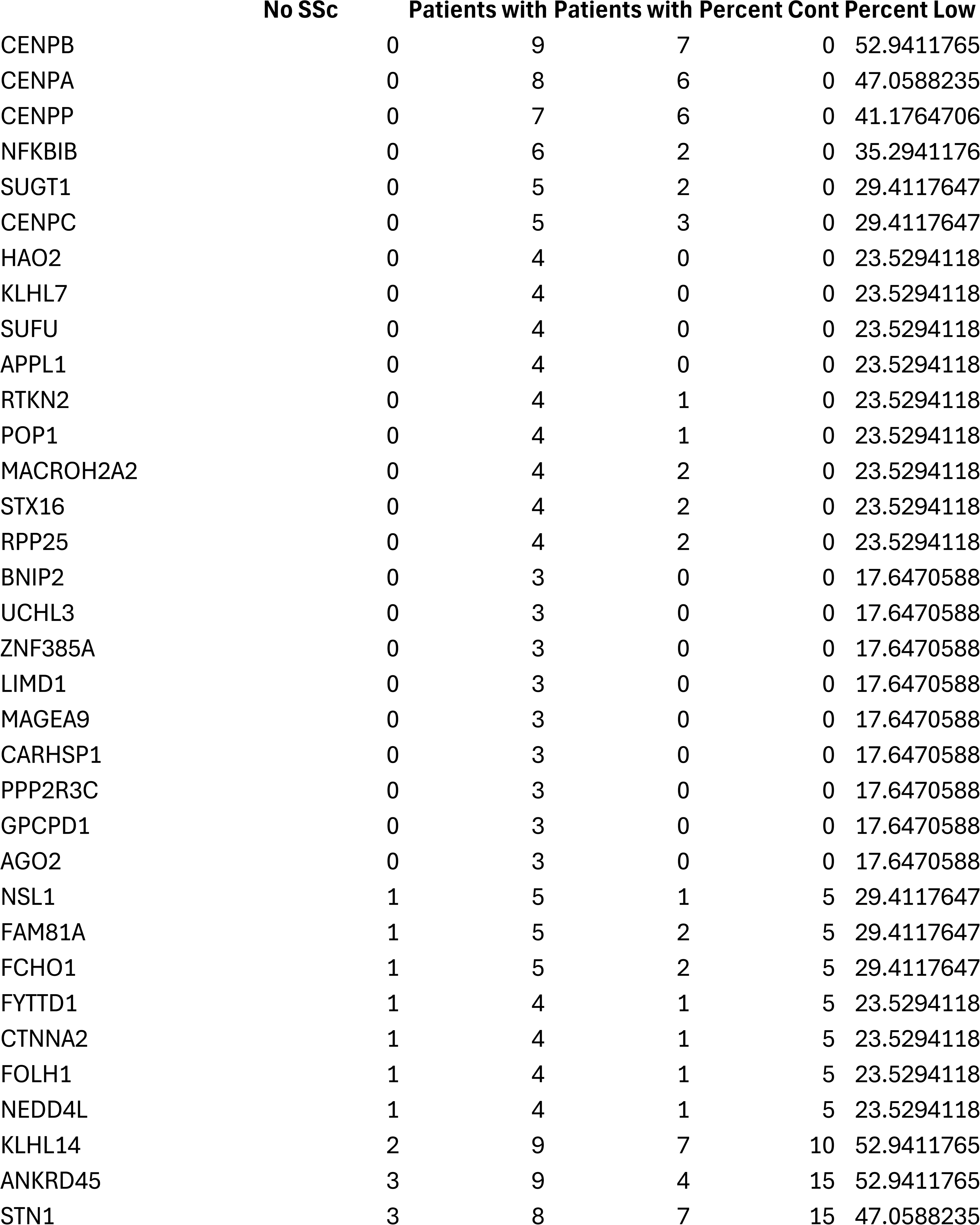
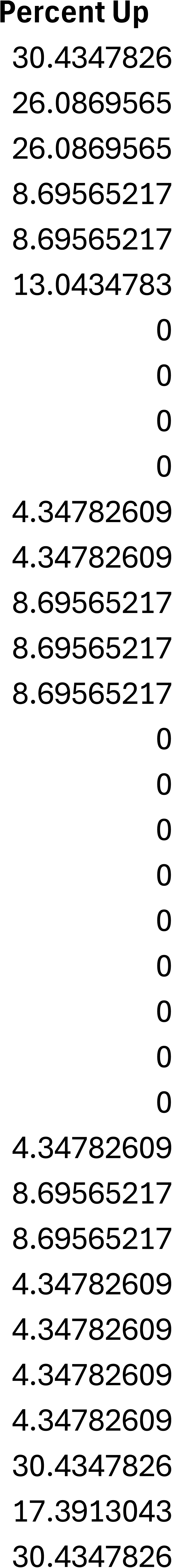
Proportion of patients with upper GI–predominant SSc, lower GI–predominant SSc, and non-SSc controls exhibiting autoantibodies against antigens significantly enriched in lower GI–associated SSc compared with the other cohorts.

|  | No SSc | Patients with | Patients with | Percent Cont | Percent Low |
| --- | --- | --- | --- | --- | --- |
| CENPB | 0 | 9 | 7 | 0 | 52.9411765 |
| CENPA | 0 | 8 | 6 | 0 | 47.0588235 |
| CENPP | 0 | 7 | 6 | 0 | 41.1764706 |
| NFKBIB | 0 | 6 | 2 | 0 | 35.2941176 |
| SUGT1 | 0 | 5 | 2 | 0 | 29.4117647 |
| CENPC | 0 | 5 | 3 | 0 | 29.4117647 |
| HAO2 | 0 | 4 | 0 | 0 | 23.5294118 |
| KLHL7 | 0 | 4 | 0 | 0 | 23.5294118 |
| SUFU | 0 | 4 | 0 | 0 | 23.5294118 |
| APPL1 | 0 | 4 | 0 | 0 | 23.5294118 |
| RTKN2 | 0 | 4 | 1 | 0 | 23.5294118 |
| POP1 | 0 | 4 | 1 | 0 | 23.5294118 |
| MACROH2A2 | 0 | 4 | 2 | 0 | 23.5294118 |
| STX16 | 0 | 4 | 2 | 0 | 23.5294118 |
| RPP25 | 0 | 4 | 2 | 0 | 23.5294118 |
| BNIP2 | 0 | 3 | 0 | 0 | 17.6470588 |
| UCHL3 | 0 | 3 | 0 | 0 | 17.6470588 |
| ZNF385A | 0 | 3 | 0 | 0 | 17.6470588 |
| LIMD1 | 0 | 3 | 0 | 0 | 17.6470588 |
| MAGEA9 | 0 | 3 | 0 | 0 | 17.6470588 |
| CARHSP1 | 0 | 3 | 0 | 0 | 17.6470588 |
| PPP2R3C | 0 | 3 | 0 | 0 | 17.6470588 |
| GPCPD1 | 0 | 3 | 0 | 0 | 17.6470588 |
| AGO2 | 0 | 3 | 0 | 0 | 17.6470588 |
| NSL1 | 1 | 5 | 1 | 5 | 29.4117647 |
| FAM81A | 1 | 5 | 2 | 5 | 29.4117647 |
| FCHO1 | 1 | 5 | 2 | 5 | 29.4117647 |
| FYTTD1 | 1 | 4 | 1 | 5 | 23.5294118 |
| CTNNA2 | 1 | 4 | 1 | 5 | 23.5294118 |
| FOLH1 | 1 | 4 | 1 | 5 | 23.5294118 |
| NEDD4L | 1 | 4 | 1 | 5 | 23.5294118 |
| KLHL14 | 2 | 9 | 7 | 10 | 52.9411765 |
| ANKRD45 | 3 | 9 | 4 | 15 | 52.9411765 |
| STN1 | 3 | 8 | 7 | 15 | 47.0588235 |

### SSc-wide and control-specific autoantibody signatures

To define disease-wide and control-enriched signals, we compared controls with the combined SSc cohort. This analysis identified autoantibodies present exclusively in SSc irrespective of GI phenotype (**Fig. 4**; **Table 4**), including those targeting EFS, CD248, WTAP, GGYF1, and NDRG4. We also identified autoantibodies enriched in SSc, but present at low frequency (5%) in controls, including those targeting SPAAR and GALNTL6 (32.5%), IFNAR1 and PXYLP1 (30%), GFAP (an astrocyte and enteric glial marker, 27.5%), IFNA17, PUF60, CLEC9A, FUT10, IZUMO4, and FUT10 (each 27.5%). Conversely, a small subset of autoantibodies (HPD, MARCKSL1, LRRFIP1, ACOT12) was detected exclusively in controls, albeit only in 15% of all control patients (**Fig. 4**; **Table 4**). These findings highlight a background repertoire of immunoreactivity that may reflect non-pathogenic or potentially protective immune signatures.

**Fig. 4:**
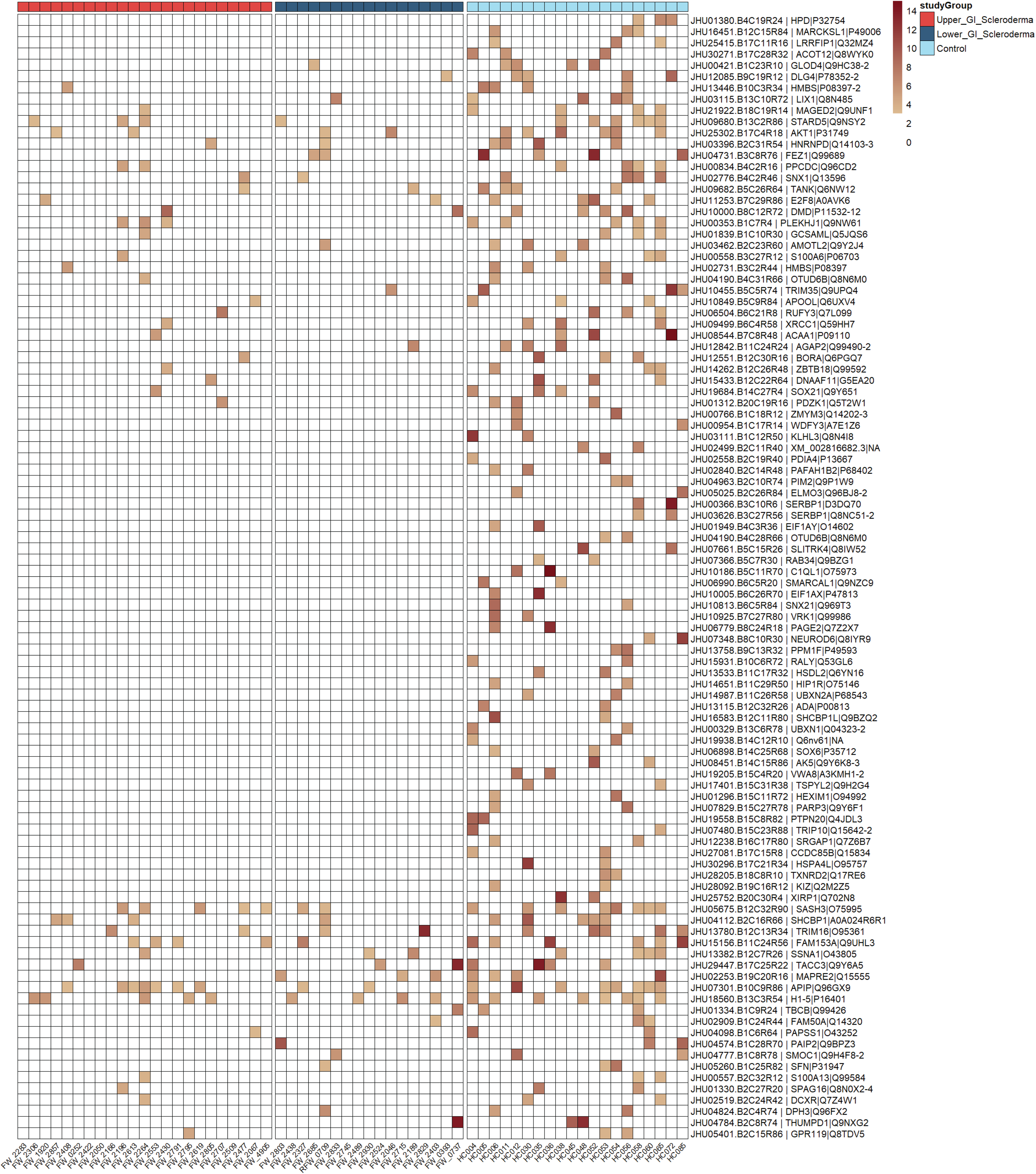
Autoantigens enriched in non⍰ISSc patients compared with SSc. Heatmap showing proteins targeted by autoantibodies significantly enriched in non-SSc individuals (light blue) relative to the combined SSc cohort (upper GI, red; lower GI, dark blue). Columns represent participants and rows represent autoantigens. Color intensity reflects relative autoantibody signal. Only autoantibodies meeting statistical significance (Fisher’s exact test, pL<L0.05; controls vs all SSc) are shown.

**Table 4:** Proportion of patients with upper GI–predominant SSc, lower GI–predominant SSc, and non-SSc controls exhibiting autoantibodies against antigens significantly enriched in patients without SSc compared with patients with SSc.

|  | No SSc | SSc | Percent Cont | Percent SSc |
| --- | --- | --- | --- | --- |
| CENPB\|P07199 | 0 | 16 | 0 | 40 |
| CENPA\|P49450 | 0 | 14 | 0 | 35 |
| CENPP\|Q6IPU0 | 0 | 13 | 0 | 32.5 |
| EFS\|O43281-2 | 0 | 13 | 0 | 32.5 |
| CENPK\|Q9BS16 | 0 | 10 | 0 | 25 |
| CD248\|Q9HCU0 | 0 | 10 | 0 | 25 |
| WTAP\|Q15007 | 0 | 9 | 0 | 22.5 |
| NDRG4\|Q9ULP0-2 | 0 | 9 | 0 | 22.5 |
| TOP1MT\|Q969P6 | 0 | 9 | 0 | 22.5 |
| GIGYF1\|O75420 | 0 | 9 | 0 | 22.5 |
| GIGYF1\|O75420 | 0 | 9 | 0 | 22.5 |
| NFKBIB\|Q15653 | 0 | 8 | 0 | 20 |
| KLHL12\|Q53G59 | 0 | 8 | 0 | 20 |
| DEFB126\|Q9BYW3 | 0 | 8 | 0 | 20 |
| CENPC\|Q03188-2 | 0 | 8 | 0 | 20 |
| LN608954.1\|NA | 1 | 14 | 5 | 35 |
| SIGLEC8\|Q9NYZ4 | 1 | 14 | 5 | 35 |
| SPAAR\|A0A1B0GVQ0 | 1 | 13 | 5 | 32.5 |
| GALNTL6\|Q49A17 | 1 | 13 | 5 | 32.5 |
| CDC25B\|P30305 | 1 | 12 | 5 | 30 |
| IFNAR1\|P17181 | 1 | 12 | 5 | 30 |
| NFU1\|Q9UMS0 | 1 | 12 | 5 | 30 |
| CD47\|Q08722 | 1 | 12 | 5 | 30 |
| PXYLP1\|Q8TE99 | 1 | 12 | 5 | 30 |
| HSPB8\|Q9UJY1 | 1 | 11 | 5 | 27.5 |
| LAT\|O43561-2 | 1 | 11 | 5 | 27.5 |
| GFAP\|P14136 | 1 | 11 | 5 | 27.5 |
| IFNA17\|P01571 | 1 | 11 | 5 | 27.5 |
| PUF60\|Q9UHX1-2 | 1 | 11 | 5 | 27.5 |
| CLEC9A\|Q6UXN8 | 1 | 11 | 5 | 27.5 |
| FUT10\|Q6P4F1-2 | 1 | 11 | 5 | 27.5 |
| IZUMO4\|Q1ZYL8 | 1 | 11 | 5 | 27.5 |
| FUT7\|Q11130 | 1 | 11 | 5 | 27.5 |
| DNAJB6\|O75190-2 | 2 | 17 | 10 | 42.5 |
| PVR\|A0A0C4DG49 | 2 | 17 | 10 | 42.5 |
| KLHL14\|Q9P2G3 | 2 | 16 | 10 | 40 |
| CBX1\|P83916 | 2 | 15 | 10 | 37.5 |
| KLRD1\|Q53ZY6 | 2 | 15 | 10 | 37.5 |
| PMEPA1\|Q969W9-3 | 3 | 24 | 15 | 60 |
| PPAT\|Q06203 | 3 | 20 | 15 | 50 |
| PIEZO1\|NA | 3 | 20 | 15 | 50 |
| CHST2\|Q9Y4C5 | 3 | 20 | 15 | 50 |
| SLC3A1\|Q07837 | 3 | 19 | 15 | 47.5 |
| TEX264\|Q9Y6I9 | 3 | 19 | 15 | 47.5 |
| GCNT2\|Q8N0V5-3 | 3 | 18 | 15 | 45 |
| SGCD\|Q92629-3 | 3 | 18 | 15 | 45 |
| P2RX4\|Q99571 | 3 | 18 | 15 | 45 |
| PACC1\|Q9H813 | 3 | 17 | 15 | 42.5 |
| FAM151A\|Q8WW52-2 | 3 | 17 | 15 | 42.5 |
| FAM20B\|O75063 | 3 | 17 | 15 | 42.5 |
| ST3GAL3\|Q11203-2 | 3 | 17 | 15 | 42.5 |
| ENPP3\|O14638 | 3 | 17 | 15 | 42.5 |
| SIGLEC5\|O15389 | 3 | 17 | 15 | 42.5 |
| HPD\|P32754 | 3 | 0 | 15 | 0 |
| MARCKSL1\|P49006 | 3 | 0 | 15 | 0 |
| LRRFIP1\|Q32MZ4 | 3 | 0 | 15 | 0 |
| ACOT12\|Q8WYK0 | 3 | 0 | 15 | 0 |
| TMEM30A\|Q9NV96 | 4 | 21 | 20 | 52.5 |
| GLOD4\|Q9HC38-2 | 4 | 1 | 20 | 2.5 |
| DLG4\|P78352-2 | 4 | 1 | 20 | 2.5 |
| HMBS\|P08397-2 | 4 | 1 | 20 | 2.5 |
| LIX1\|Q8N485 | 4 | 1 | 20 | 2.5 |
| MAGED2\|Q9UNF1 | 4 | 1 | 20 | 2.5 |
| PRPH2\|P23942 | 5 | 24 | 25 | 60 |
| ENTPD3\|O75355-2 | 6 | 26 | 30 | 65 |
| ERP27\|Q96DN0 | 8 | 28 | 40 | 70 |
| SPN\|P16150 | 9 | 32 | 45 | 80 |
| PMEPA1\|Q969W9-2 | 9 | 32 | 45 | 80 |
| ARMH4\|Q86TY3 | 14 | 38 | 70 | 95 |

Table 4: Clinical features of the 4 Systemic sclerosis cohorts
| Variable | $\alpha$ | $\beta$ | $\gamma$ | $\delta$ | p-value |
| --- | --- | --- | --- | --- | --- |
| N | 14 | 5 | 5 | 16 |  |
| Age*, years, mean (SD) | 50 (14) | 47 (10) | 60 (7) | 57 (10) | 0.12 |
| Male, n (%) | 4 (29) | 0 (0) | 0 (0) | 2 (12) | 0.43 |
| Disease duration*, years (median, IQR) | 8 (5, 21) | 8 (5, 31) | 17 (10,18) | 12 (5, 23) | 0.90 |
| Race |  |  |  |  |  |
| White | 9 (64) | 4 (80) | 5 (100) | 15 (94) |  |
| Black | 4 (29) | 1 (20) | 0 (0) | 1 (6) |  |
| Other | 1 (8) | 0 (0) | 0 (0) | 0 (0) | 0.40 |
| Diffuse cutaneous, n (%) | 7 (54) | 0 (0) | 2 (40) | 0 (0) | **0.001** |
| Ever smoker, n (%) | 3 (21) | 2 (40) | 2 (40) | 6 (40) | 0.68 |
| Organ severity and involvement |  |  |  |  |  |
| Significant Raynaud<sup>†</sup>s<sup>^</sup> ($\geq 2$), n (%) | 5 (36) | 4 (80) | 0 (0) | 5 (33) | **0.07** |
| Cardiac involvement<sup>^</sup> (Medsger $\geq 1$), n (%) | 2 (25) | 0 (0) | 0 (0) | 1 (14) | 0.67 |
| Significant GI<sup>^</sup> (Medsger $\geq 2$), n (%) | 12 (86) | 3 (60) | 4 (80) | 9 (56) | 0.31 |
| Muscle weakness<sup>^</sup> (Medsger $\geq 2$), n (%) | 0 (0) | 0 (0) | 0 (0) | 2 (13) | 0.73 |
| Pulmonary Fibrosis, n (%) | 11 (92) | 2 (50) | 3 (60) | 2 (20) | **0.004** |
| Cardiopulmonary testing |  |  |  |  |  |
| Minimum FVC, mean (SD) | 67 (20) | 89 (16) | 85 (8) | 91 (15) | **0.004** |
| Minimum DLCO | 45 (19) | 67 (16) | 62 (21) | 60 (25) | 0.15 |
| Maximum RVSP, mmHg (median, IQR) | 39 (38, 44) | 31 (23, 32) | 28 (25, 30) | 28 (26, 43) | 0.10 |
| Previously Scored Autoantibodies |  |  |  |  |  |
| Topoisomerase-1 (Scl70) | 2 (20) | 3 (60) | 1 (20) | 0 (0) | **0.03** |
| RNA polymerase 3 | 1 (10) | 0 (0) | 0 (0) | 0 (0) | 0.65 |
| Centromere (A or B) | 0 (0) | 0 (0) | 0 (0) | 10 (91) | **<0.001** |
| RNPC3 (U11/U12) | 3 (21) | 1 (20) | 0 (0) | 0 (0%) | 0.16 |
| U1RNP | 4 (29) | 0 (0) | 0 (0) | 2 (12) | 0.43 |
| U3RNP (Fibrillarin) | 1 (7) | 0 (0) | 0 (0) | 2 (12) | 1.00 |
| Ku | 0 (0) | 0 (0) | 0 (0) | 1 (9) | 1.00 |
| Th/To | 0 (0) | 0 (0) | 1 (20) | 0 (0) | 0.32 |
| Gephyrin | 0 (0) | 0 (0) | 0 (0) | 2 (12) | 0.71 |
| Ro52 | 6 (43) | 2 (40) | 1 (20) | 5 (31) | 0.86 |
| Ago (1 or 2) | 2 (14) | 0 | 0 | 1 (6) | 0.83 |

\*Age and disease duration are from the time of the WGT; <sup>^</sup>=Medsger score; FVC = forced vital capacity, DLCO = Diffusing capacity, RVSP = estimated right ventricular systolic pressure on echo

### Serology-driven clustering revealed biologically distinct and clinically relevant subgroups within SSc

Although several autoantibodies showed clear disease associations, no single autoantibody was exclusive to a given clinical phenotype. This observation suggests that stratification based solely by clinical manifestations, such as upper or lower GI involvement, may obscure underlying serological heterogeneity and limit accurate disease classification. To address this, we performed a bottom-up unsupervised clustering analysis using autoantibodies enriched in patients with upper GI disease relative to controls. This approach identified three clusters (A–C) (**Fig. 5**), with corresponding seromic patterns (i.e. autoantibodies that together were enriched in each of the three clusters) summarized in **Supplementary Data 1**. Cluster A was the smallest, comprising 6 patients (4 with lower GI disease and 2 with upper GI disease, with no controls). Cluster C included 17 patients (12 upper GI, 3 lower GI, and 2 controls), whereas Cluster B lacked distinctive seromic enrichment, suggesting that clustering derived from upper GI-associated features alone did not yield clearly discriminative or clinically informative subgroups.

**Fig. 5:**
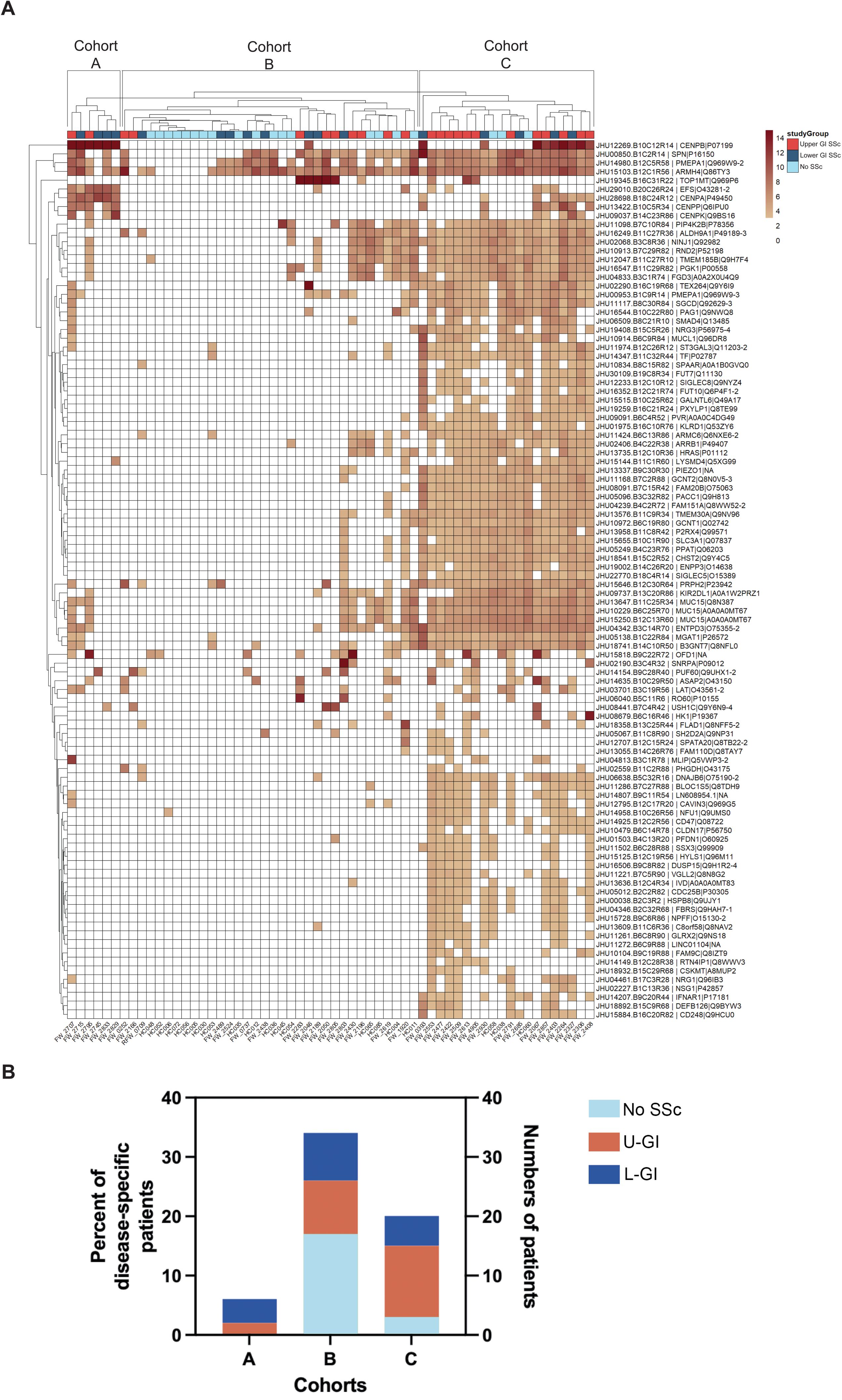
Serology-defined clustering based on upper GI-enriched autoantibodies. (A) Heatmap and hierarchical clustering identifying three patient clusters (A–C) based on autoantibodies significantly enriched in upper GI SSc relative to controls (Fisher’s exact test, pL<L0.05). Rows indicate autoantigens; columns represent individual participants. Color intensity denotes relative autoantibody signal. (B) Distribution of clinical subgroups within each cluster, including upper GI SSc (red), lower GI SSc (dark blue), and controls (light blue). Cohort A is enriched for SSc without controls; cohort B contains the highest proportion of controls; cohort C is enriched for patients with lower GI disease.

We therefore performed a complementary analysis using autoantibodies enriched in patients with lower GI disease. This independent approach identified four clusters (α, β, γ, and δ) (**Fig. 6**), with integrated clinical and serological features summarized in **Table 5** and detailed seromic patterns provided in **Supplementary Data 2**. Cluster composition demonstrated marked differences in disease distribution. Cluster α consisted of 18 patients (12 upper GI, 2 lower GI, and 4 controls) and was enriched in upper GI disease (66.7%). Cluster γ, the smallest group, included 6 patients (5 lower GI, 1 control) and was strongly enriched for lower GI disease (83/3%). Cluster δ included 17 patients (8 upper GI, 8 lower GI, and 1 control). Cluster composition revealed clear differences in disease distribution. Cluster α was dominated by upper GI disease patients (66.67%), whereas cluster γ was dominated by lower GI disease patients (83.34%). Cluster δ showed a balanced representation of upper and lower GI disease (47.05% each), while cluster β was composed primarily of controls (73.68%). This parallel analytical approach enabled independent derivation of phenotype-enriched feature sets, allowing assessment of whether consistent clustering structures emerged across complementary analyses.

**Fig. 6:**
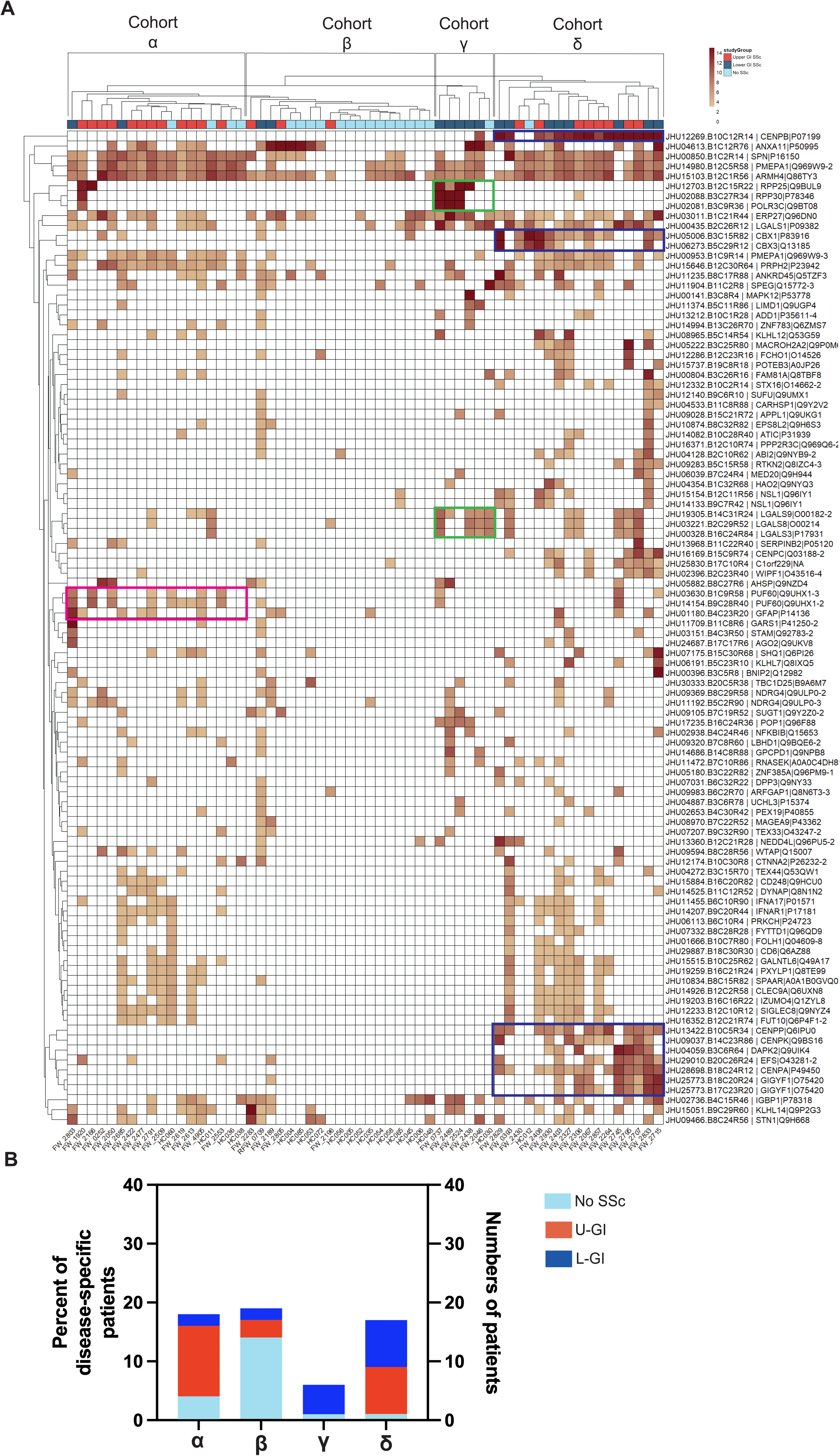
Serology-defined clustering based on lower GI–enriched autoantibodies. (A) Heatmap and hierarchical clustering identifying four patient clusters (α–δ) based on autoantibodies significantly enriched in lower GI SSc relative to controls (Fisher’s exact test, pL<L0.05). Rows indicate autoantigens; columns represent participants. Color intensity denotes relative autoantibody signal. Colored annotations highlight cluster-defining autoantigens: cohort α (PUF60, GFAP); cohort γ (RPP25, RPP30, POLR3C); cohort δ (CENPB, CBX1, CBX3, CENPP, CENPK, DAPK2, EFS, CENPA, GIGYF1). Cohort β lacks defining autoantigens. (B) Clinical composition of clusters. Cohort α is enriched for upper GI disease, cohort β for controls, cohort γ for lower GI disease, and cohort δ includes a balanced representation of upper and lower GI SSc.

**Table 5:** Clinical features of the four SSc cohorts α, β, γ, and δ.

| Variable | $\alpha$ | $\beta$ | $\gamma$ | $\delta$ | p-value |
| --- | --- | --- | --- | --- | --- |
| Esophageal at 10s,<br>median (IQR) | 82 (65, 85) | 79 (70, 85) | 90 (88, 91) | 81 (28, 90) | 0.14 |
| Solid Gastric at 4hr,<br>median (IQR) | 96 (95, 97) | 97 (96, 97) | 97 (94, 98) | 97 (95, 98) | 0.92 |
| Small Bowel at 6hr,<br>median (IQR) | 69 (62, 82) | 68 (64, 85) | 67 (66, 76) | 65 (53, 77) | 0.70 |
| Large Bowel at 72hr,<br>median (IQR) | 91 (78, 93) | 44 (0, 90) | 0 (0, 0) | 71 (19, 86) | **0.005** |

For reference: Delayed Esophageal transit at 10s is <83%, Delayed solid emptying is less than 90% at 4 hours. Delayed small bowel transit is <49% at 6 hours. Delayed large bowel transit is <67% by 72 hours.

The seromic and clinical GI patterns for the 4 clusters can be defined as the following

- *<u>Cluster</u>* <u>δ</u>: 17 patients (8 upper GI, 8 lower GI, and 1 control), suggesting equal representation of upper and lower GI disease incidence; defined by autoantibodies against CBX1, CBX3, CENP (Centromere)-family antigens, DAPK2, EFS, and GIGYF1.
- *<u>Cluster</u>* <u>γ</u>: 6 patients (0 upper GI, 5 lower GI, and 1 control), suggesting significant lower GI disease incidence; harbors autoantibodies against RPP25, RPP30, POLR3C and LGALS9, while not harboring Cohort δ-specific autoantibodies. RPP25 (Ribonuclease P/MRP Subunit P25) is a 25 kDa component of the ThTo complex^39^, and RPP30 (Ribonuclease P/MRP Subunit P30), is a 30 kDa component of the same complex^40^, suggesting that anti-ThTo antibodies are a defining seromic pattern of this cluster
- *<u>Cluster</u>* <u>α</u>: 18 patients (12 with upper GI, 2 with lower GI, and 4 controls), suggesting significant upper GI disease incidence; harbors autoantibodies against PUF60 and GFAP, while not harboring Cohort δ-specific autoantibodies.
- Finally, *<u>Cluster</u>* <u>β</u>: 19 patients (3 upper GI, 2 lower GI, and 14 controls), highest proportion of control patients, and has no defining seromic pattern.

We next examined whether serology-defined clusters corresponded to distinct clinical phenotypes spanning vascular, cardiopulmonary, GI, and autonomic domains (**Tables 5–7**) and found that each cluster was associated with a reproducible and clinically interpretable pattern of disease. Cluster α exhibited a multisystem *fibrotic phenotype* with prominent pulmonary involvement, including the highest rates of pulmonary fibrosis (92%, p = 0.004) and severe GI disease (86%), along with the lowest lung function (FVC 67% ± 20). SSc patients in Cluster β were characterized by a *vasculopathic/autonomic phenotype* with preserved cardiopulmonary function but prominent GI and autonomic dysfunction. Cluster γ included the oldest patients with the longest disease duration, and this cluster showed *a lower GI–predominant* phenotype with severe colonic hypomotility (0% large bowel emptying at 72 h; p = 0.005) despite preserved pulmonary function, and finally, Cluster δ represented a *limited cutaneous phenotype* with minimal pulmonary fibrosis (20%; p < 0.001) and preserved lung function.

**Table 6:** Percent GI emptying by region in the four SSc cohorts α, β, γ, and δ.

**Table 7:** Symptoms of autonomic dysfunction (COMPASS-31 scores) in the four SSc cohorts α, β, γ, and δ.

Importantly, autoantibodies against TiSSc1/2 were detected in clusters α (35.71%) and δ (56.25%) but were not significantly enriched in a single cluster (p = 0.29), indicating that TiSSc1/2 reactivity spans multiple serological contexts rather than defining a unique disease subset.

These findings demonstrate that serological clustering corresponds to clinically meaningful disease phenotypes across multiple organ systems.

### Novel serology provides superior discrimination of GI phenotypes compared to canonical autoantibodies

We evaluated whether established SSc autoantibody profiles could be recapitulated using HuProt. Known seropositivity for AGO1, centromere, and Scl⍰70 was accurately detected (**Supplementary Fig. 7A**), with limited power for rarer specificities. However, canonical autoantibodies provided limited discrimination between upper and lower GI disease (**Supplementary Fig. 7B**). Many were either infrequent (e.g., anti–Th/To) or similarly distributed across GI phenotypes (e.g., anti⍰CENPB). In contrast, newly identified autoantibodies demonstrated clear subtype specificity; for example, anti⍰TiSSc1/2 antibodies were detected in approximately 50% of patients with upper GI disease compared with ∼10% of those with lower GI disease. Together, these findings demonstrate that proteome-wide serological profiling reveals previously unrecognized, biologically structured heterogeneity in SSc-associated GI disease, enabling identification of clinically meaningful patient subgroups that extend beyond traditional anatomy-based classification.

## Discussion

In this study, we applied unbiased, proteome-wide serological profiling to patients with SSc and GI involvement to test the hypothesis that distinct patterns of autoimmunity underlie clinically divergent GI phenotypes. We used parallel comparisons to identify phenotype-enriched signals that might be diluted in multi-group analyses within a high-dimensional feature space, and to assess whether independently derived autoantibody sets converged on consistent clustering structures. The convergence observed across these complementary approaches supports the robustness of the identified serological subgroups. These findings directly address the central premise that the heterogeneity of SSc-associated GI disease reflects underlying biological differences rather than purely anatomic variation. By integrating high-dimensional autoantibody data with detailed clinical phenotyping, we demonstrate that the structure of the humoral immune response captures clinically meaningful disease subsets characterized by distinct patterns of organ involvement, physiological impairment, and disease burden.

A central finding of this work is the identification of novel autoantibody repertoires that discriminate between upper and lower GI disease. These signatures extend beyond canonical SSc-associated autoantibodies, which in our cohort were either infrequently detected or failed to distinguish clinically relevant GI subtypes (**Supplementary Fig. 7B**). In contrast, newly identified autoantibodies, including those targeting previously unannotated or poorly characterized proteins, demonstrated clear phenotype-specific enrichment. Building on our prior work, we further validated autoantibodies against AGO2 in SSc and demonstrated that both their presence and abundance correlate with constipation incidence and severity, with specificity for patients with lower GI involvement^41^. We observe autoantibodies against PUF60, which have been previously described in primary Sjögren’s syndrome and other rheumatologic diseases^42^. This study extends the relevance of anti-PUF60 autoantibodies to a distinct subset of patients with SSc, linking this immune signature to a severe, multi-system phenotype. Furthermore, prior studies have associated hepatitis B infection with SSc^43^. Together, our findings indicate that the serological landscape of SSc-associated GI disease is broader than currently appreciated and captures dimensions of disease biology not reflected in standard clinical or serological frameworks.

Among these discoveries, we identified LN608954.1 (designated TiSSc1/2) as a previously unrecognized autoantigen enriched in upper GI disease. While its biological function remains incompletely defined, we provide initial evidence that TiSSc is expressed in human cells, localizes predominantly to the cytoplasm, and is responsive to cellular stress (**Fig. 2; Supplementary Fig. 6**). Anti⍰TiSSc1/2 reactivity was associated with delayed esophageal transit, suggesting a relationship with clinically meaningful dysfunction. However, TiSSc1/2 antibodies were not restricted to a single serological cluster, and their functional significance remains uncertain. Accordingly, despite its relatively high prevalence, the designation of TiSSc as a disease-relevant autoantigen warrants further investigation, particularly into the functional roles of these proteins and the mechanisms by which their targeting may contribute to esophageal dysmotility.

The presence of autoantibodies in control individuals and the detection of higher-prevalence signals across groups warrant careful interpretation. Low-level autoreactivity is a recognized feature of the normal immune repertoire and may reflect physiological immune surveillance, cross-reactivity, or prior environmental and infectious exposures rather than disease-specific processes. These factors may contribute to shared or background signals across SSc and control samples. Importantly, disease associations in this study were defined by patterns of enrichment rather than binary presence alone. Nonetheless, distinguishing disease-relevant signals from background immunoreactivity will require validation in independent cohorts and, where feasible, confirmation using orthogonal assays.

Beyond individual autoantibodies, our analyses demonstrate that proteome-wide serological profiling enables clinically meaningful stratification of SSc patients with GI involvement. Clustering based on shared autoantibody patterns identified reproducible subgroups with distinct organ involvement, physiological profiles, and clinical features (**Figs. 5–6**; **Tables 5–7**). These serology-defined clusters aligned with established disease phenotypes, including centromere-associated limited cutaneous disease and Th/To-related lower GI involvement, providing biological plausibility and internal validation. Although derived from a modest cohort and therefore requiring cautious interpretation, the convergence of clustering across independent analytical approaches and its concordance with known disease biology suggest that these subgroups reflect reproducible structure rather than purely data-driven artifacts.

Collectively, these findings extend beyond the identification of individual autoantibodies to define a higher-order organization of the humoral immune response in SSc. The observed clustering patterns are consistent with the concept that region-specific GI dysfunction may arise from distinct underlying biological processes, potentially involving differential contributions of neural, stromal, or immune pathways, although direct mechanistic links remain to be established. Together, these data suggest that SSc GI disease is not a single entity but rather comprises biologically distinct subtypes defined by coordinated immune signatures, supporting a shift from anatomy-based to immune-informed classification.

From a patient perspective, GI involvement in SSc is highly heterogeneous and difficult to predict, with substantial and often unpredictable variability in onset, severity, and progression. The inability to identify which patients will develop clinically significant GI dysfunction represents a major unmet need, particularly given the associated risks of malnutrition, recurrent hospitalization, and increased mortality. The absence of reliable biomarkers further limits risk stratification and clinical decision-making. In this context, our findings suggest that serology-based stratification may provide a complementary framework to improve patient phenotyping and risk assessment, although these applications remain investigational.

These findings also have implications for biomarker development and clinical trial design. The identification of subgroup-specific autoantibody signatures supports the feasibility of developing blood-based tools for disease stratification; however, their clinical utility will depend on independent validation, reproducibility, and demonstration of added value beyond existing measures. Similarly, the substantial heterogeneity observed across serological and clinical features suggests that treating SSc GI disease as a single entity may dilute therapeutic signals. Serology-informed stratification may therefore enable enrichment of more biologically homogeneous trial populations and improve the detection of treatment effects, although this approach requires prospective evaluation.

Several limitations warrant consideration when interpreting these findings. First, this study relies on comprehensive seromic analyses performed in a relatively limited number of patient samples. Given the high-dimensional nature of proteome-wide screening, the absence of formal correction for multiple comparisons introduces the possibility of false-positive associations. Although the consistency of findings enrichment across complementary analytical frameworks and their convergence with established disease biology mitigates this concern, candidate autoantibodies should be interpreted cautiously and prioritized for independent validation. Second, while the sample size is sufficient to detect robust signals, the resource-intensive nature of these analyses limits the resolution of less prevalent serological patterns. To minimize overinterpretation, we did not perform subgroup analyses within the identified clusters, although the observed data suggest that potentially informative, cluster-specific seromic signatures may exist. In addition, although prior work has demonstrated that SSc autoantibodies targeting intracellular antigens can be functionally relevant, the pathogenic or mechanistic roles of the novel autoantibodies identified here remain to be established. Finally, this study was conducted within a single-center cohort enriched for patients with isolated upper or lower GI disease involvement, representing phenotypic extremes. As such, these findings may not fully generalize to the broader SSc population, in which mixed or evolving GI phenotypes are more common. Our study establishes a foundation for future work demonstrating that comprehensive seromic profiling in larger, multi-center cohorts of patients with SSc, encompassing the full spectrum of disease, is needed to more fully define biological and clinical heterogeneity. Such efforts will be essential for the development of refined clinical biomarkers and the identification of targeted therapeutic strategies. Despite these limitations, several features support the internal robustness of our findings, including the reproducibility of autoantibody enrichment across analytical approaches, convergence of clustering derived from independently defined feature sets, and concordance with well-established clinical phenotypes. Collectively, these observations strengthen confidence in the underlying structure of the data while underscoring the need for prospective validation. Looking forward, our findings support a shift toward a systems-level, immune-informed framework for disease classification of SSc.

In summary, our findings uncover previously unrecognized serological diversity in SSc-associated GI disease and demonstrate that autoantibody profiles can stratify patients into clinically meaningful subgroups. By moving beyond conventional classification toward immune-informed stratification, this work establishes a foundation for improved patient phenotyping, biomarker development, and targeted therapeutic strategies in SSc. Collectively, these data support a shift from anatomy-based to serology-defined classification of SSc GI disease.

## Acknowledgements

ZM was supported by grant funding from Rheumatology Research Foundation, the Scleroderma Research Foundation, and the NIH/NIAMS R01AR081382 grant. SK was supported by grant funding from Harvard Digestive Disease Core and NIH/NIA grant R01AG066768. AS was supported by NIH/NIAMS K24 AR080217. The Johns Hopkins Scleroderma Center Research Registry is supported by NIH/NIAMS P30 AR070254, the Johns Hopkins in Health initiative, the Donald B. and Dorothy L. Stabler Foundation, the Chresanthe Staurulakis Memorial Fund, and the Sara and Alex Othon Research Fund.

## Supplementary Data

**Supplementary Data 1. Seromic patterns of three Clusters generated by comparing autoantibodies significantly enriched in upper GI disease patients compared to controls.**

**Supplementary Data 2. Seromic patterns of four Clusters generated by comparing autoantibodies significantly enriched in lower GI disease patients compared to controls.**

### Supplementary Figures

**Supplementary Fig 1. Genomic organization of TiSSc1 and TiSSc2.** (A) Exonic and 3′ untranslated region (UTR) structure of TiSSc1 and TiSSc2 within the human genome; translated sequences are highlighted in red. (B) Genomic mapping shows that TiSSc1 and TiSSc2 share exons 1–2, whereas TiSSc2 exons 3–4 are located within intron 2 of the TiSSc1 locus. Let-7a/b microRNA sequences reside within the 3′UTR of both transcripts.

**Supplementary Fig 2. Primate homology of TiSSc sequences**. Nucleotide BLAST of TiSSc1/2 exon sequences identifies homologous regions within the genome of pygmy chimpanzee (Pan paniscus).

**Supplementary Fig 3. Evolutionary conservation of TiSSc2**. Alignment of the TiSSc2 open reading frame (first 222 bp) across placental mammals demonstrates conservation within Old World monkeys but not in other species.

**Supplementary Fig 4. Evolutionary origin of TiSSc1/2**. Comparative genomic analysis shows conservation across hominoids and select cercopithecoids, with absence in New World monkeys, suggesting emergence in Old World primates >25 million years ago.

**Supplementary Fig 5 Proteomic evidence for TiSSc1 expression**. Publicly available mass spectrometry datasets (OpenProt) demonstrate detection of TiSSc1-derived peptides across multiple human tissues.

**Supplementary Fig 6. Stress-induced modulation of TiSSc2**. LX-2 cells exposed to hyperthermia show (A) increased TiSSc2 protein abundance (mean fluorescence intensity), without corresponding changes in (B, C) let-7a/b transcript levels. These findings suggest that stress-induced TiSSc2 upregulation is not driven by increased MIRLET7BHG transcription.

**Supplementary Fig. 7. Validation and comparison of autoantibody profiles**. (A) HuProt-based validation of canonical autoantibodies (Ago1, centromere, Scl-70, Ro52/TRIM21), demonstrating concordance with established seropositivity profiles. (B) Distribution of canonical versus newly identified autoantibodies across SSc-GI subgroups. Canonical antibodies show limited discriminatory capacity, whereas novel autoantibodies, including anti-TiSSc1/2, demonstrate subtype-specific enrichment and improved separation of upper versus lower GI phenotypes.

**Supplementary Table 1: Clinical and physiological features in SSc patients with and without TiSSc seropositivity.**

