## Supplementary figures and images for "Proteome-wide serology reveals immune-defined subtypes of gastrointestinal disease in systemic sclerosis"

### Supplementary Figure 1

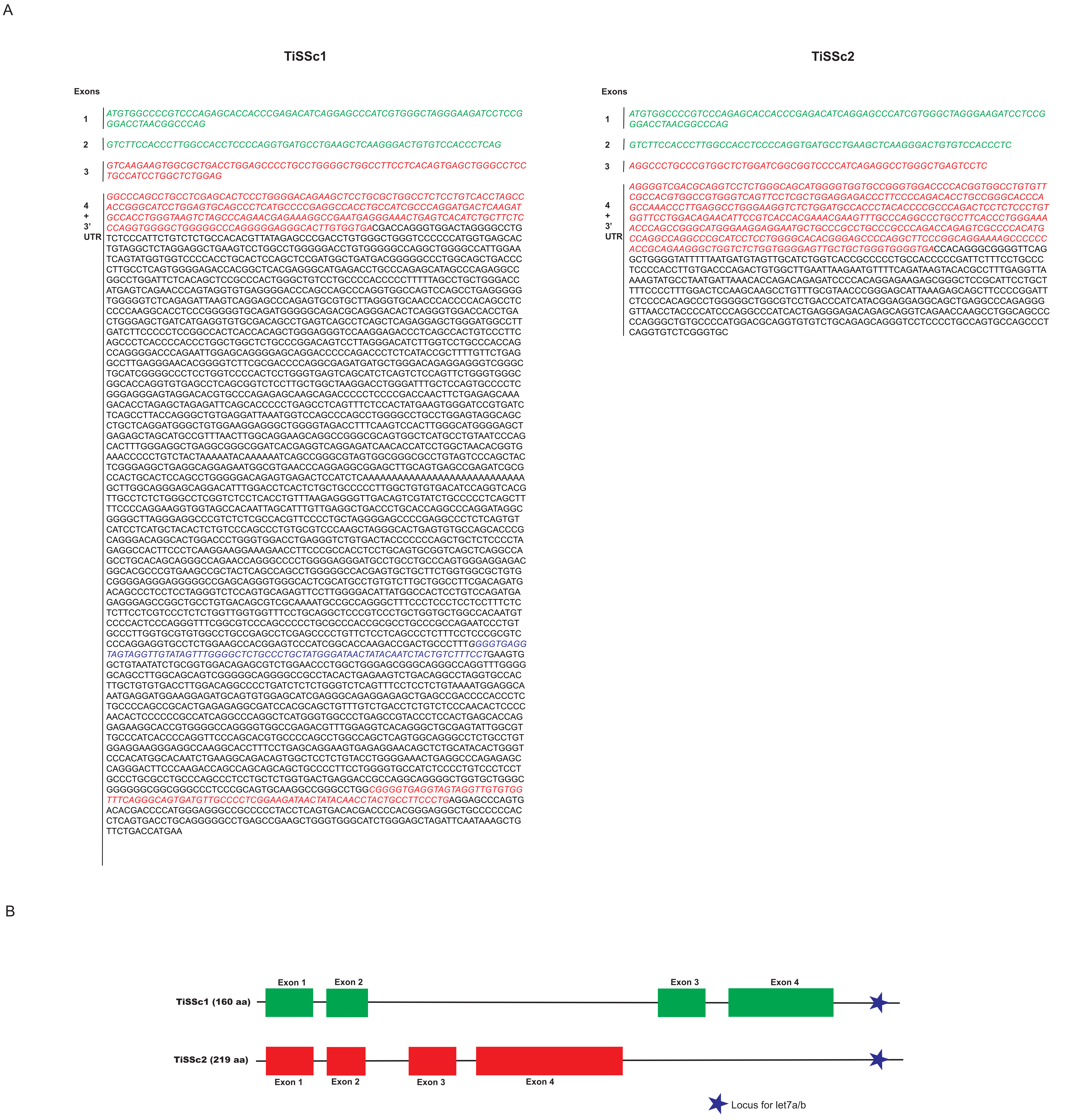

### Supplementary Figure 3

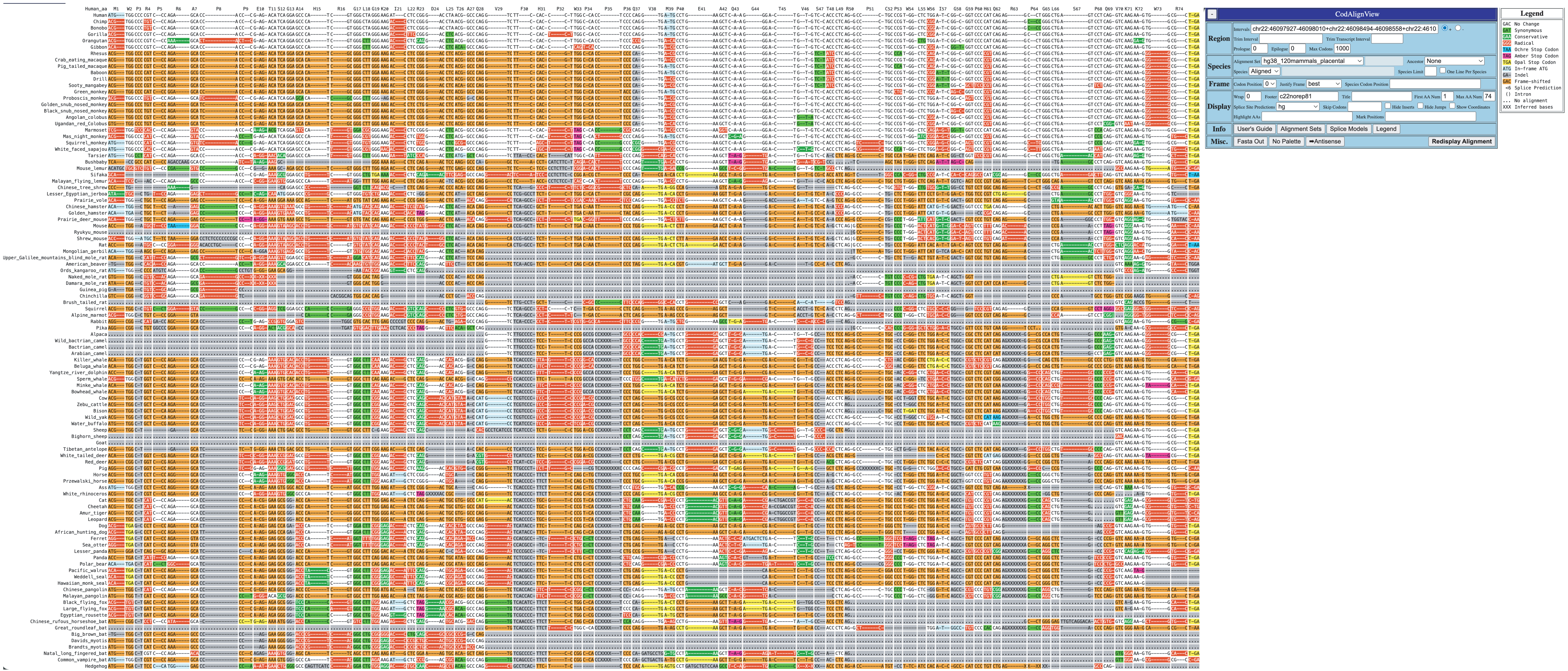

### Supplementary Figure 4

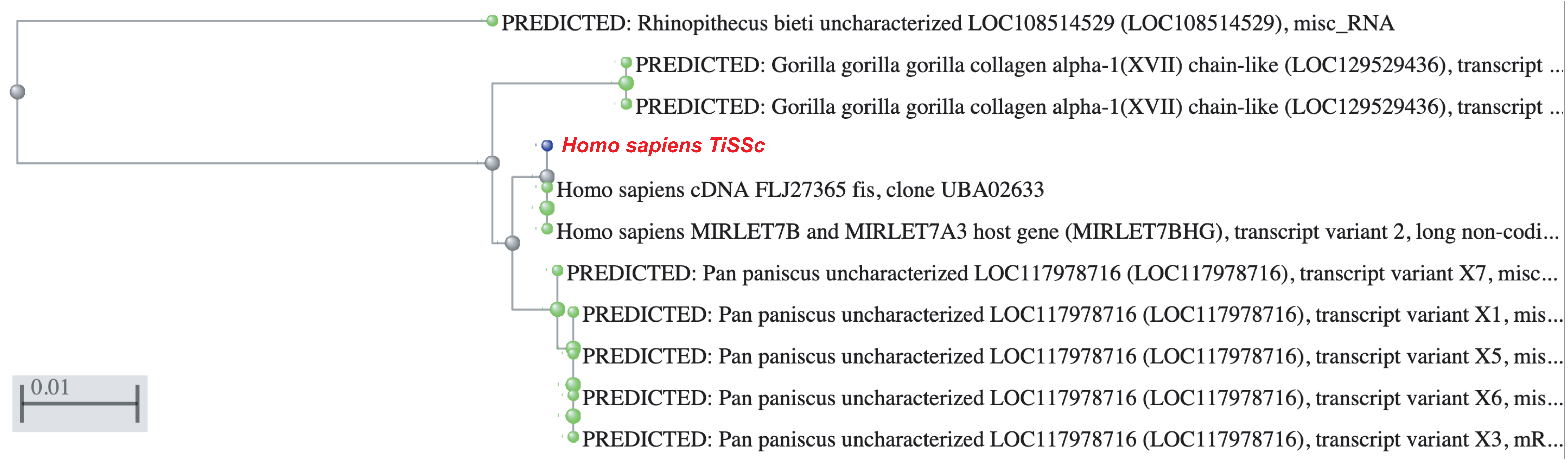

### Supplementary Figure 6

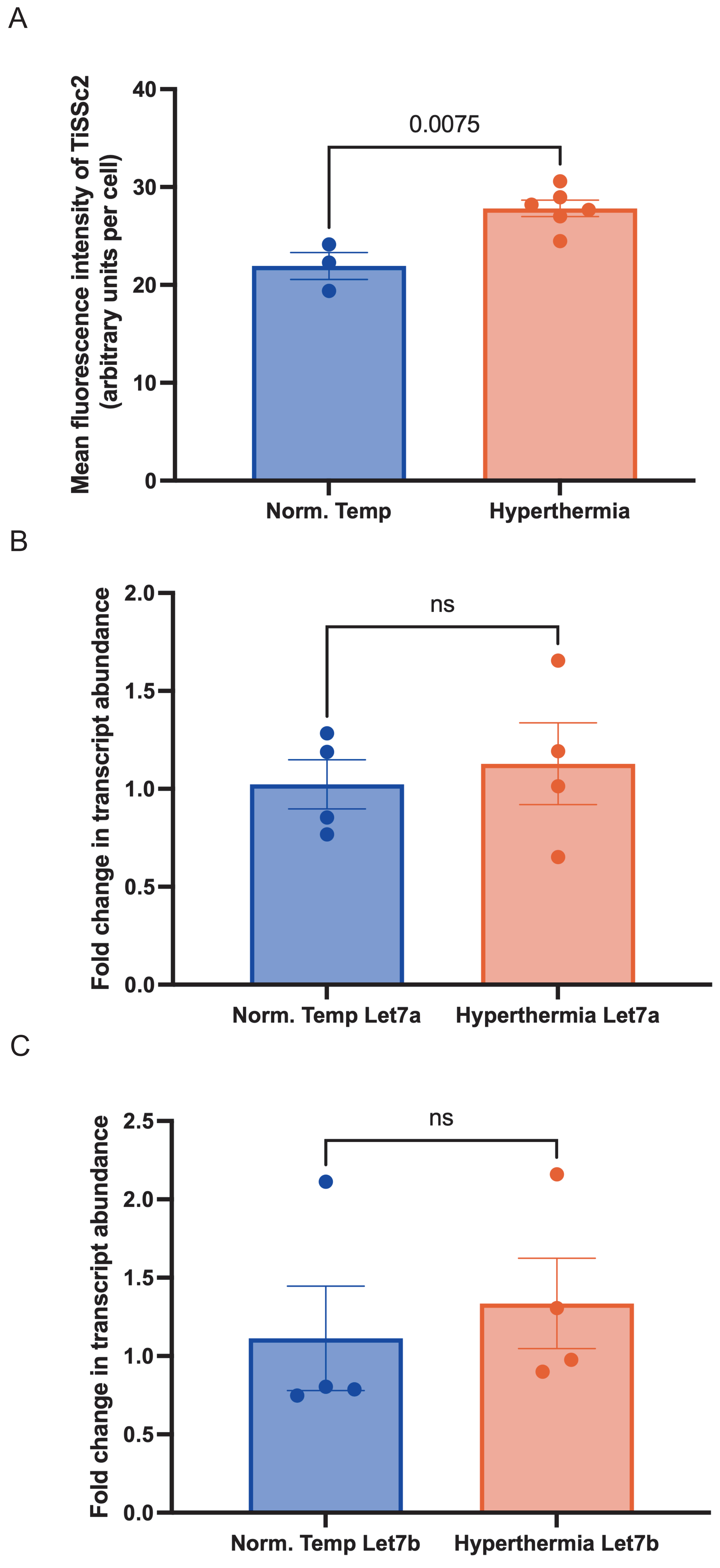
