## Supplementary Figure 2 for "Proteome-wide serology reveals immune-defined subtypes of gastrointestinal disease in systemic sclerosis"

### BLAST against *Pan paniscus* isolate Mhudiblu (Carbone #601152) 000362F_545157_qpd, whole genome shotgun sequence; Sequence ID: SSBP03005811.1

#### TiSSc1

**Exon 1**

```
Query  3       GTGGCCCCGTCCCAGAGCACCACCCGAGACATCAGGAGCCCATCGTGGGCTAGGGAAGAT  62
               ||||||| |||||||||||||||||||||||||||||||||||||||||||||||||||| 
Sbjct  174548  GTGGCCCTGTCCCAGAGCACCACCCGAGACATCAGGAGCCCATCGTGGGCTAGGGAAGAC  174607

Query  63      CCTCCGGGACCTAACGGCCCAGGT  86
               ||||||||||||||||||||||||
Sbjct  174608  CCTCCGGGACCTAACGGCCCAGGT  174631
```

**Exon 2**

```
Query  81      CCAGGTCTTCCACCCTTGGCCACCTCCCCAGGTGATGCCTGAAGCTCAAGGGACTGTGTC  140
               ||||||||||||||||||||||||||||||||||||||||||||||||||||||||||||
Sbjct  175110  CCAGGTCTTCCACCCTTGGCCACCTCCCCAGGTGATGCCTGAAGCTCAAGGGACTGTGTC  175169

Query  141     CACCCTCAGGTCAGT  155
               |||||||||||||||
Sbjct  175170  CACCCTCAGGTCAGT  175184
```

**Exon 3**

```
Query  154     GTCAAGAAGTGGCGCTGACCTGGAGCCCCTGCCTGGGGCTGGCCTTCCTCACAGTGAGCT  213
               ||||||||||||||||||||||||||||||||||||||||||||||||||||||||||||
Sbjct  185979  GTCAAGAAGTGGCGCTGACCTGGAGCCCCTGCCTGGGGCTGGCCTTCCTCACAGTGAGCT  186038

Query  214     GGGCCTCCTGCCATCCTGGCTCTGGAGG  241
               ||||||||||||||||||||||||||||
Sbjct  186039  GGGCCTCCTGCCATCCTGGCTCTGGAGG  186066
```

**Exon 4**

```
Query  234     TCTGGAGGGCCCAGCCTGCCTCGAGCACTCCCTGGGGACAGAAGCTCCTGCGCTGGCCTC  293
               |||| |||||||||||||||||||||||||||||||||||||||||||||| ||||||||
Sbjct  186473  TCTGCAGGGCCCAGCCTGCCTCGAGCACTCCCTGGGGACAGAAGCTCCTGCACTGGCCTC  186532

Query  294     TCCTGTCACCTAGCCACCGGGCATCCTGGAGTGCAGCCCTCATGCCCCGAGGCCACCTGC  353
               |||||||||||||||||||||||| |||||||||||||||||||||||||||||||||||
Sbjct  186533  TCCTGTCACCTAGCCACCGGGCATTCTGGAGTGCAGCCCTCATGCCCCGAGGCCACCTGC  186592

Query  354     CATCGCCCAGGATGACTCAAGATGCCACCTGGGTAAGTCTAGCCCAGAACGAGAAAGGCC  413
               ||||||||||||||||||||||||||||||||||||||||||||||||||||||||||||
Sbjct  186593  CATCGCCCAGGATGACTCAAGATGCCACCTGGGTAAGTCTAGCCCAGAACGAGAAAGGCC  186652

Query  414     GAATGAGGGAAACTGAGTCACATCTGCTTCTCCCAGGTGGGGCTGGGGGCCCAGGGGGAG  473
               ||||||||||||||||||||||||||||||||||||||||||||||||||||||||||||
Sbjct  186653  GAATGAGGGAAACTGAGTCACATCTGCTTCTCCCAGGTGGGGCTGGGGGCCCAGGGGGAG  186712

Query  474     GGCACTTGTGGTGA  487
               ||||||||||||||
Sbjct  186713  GGCACTTGTGGTGA  186726
```

#### TiSSc2

```
Query  3       GTGGCCCCGTCCCAGAGCACCACCCGAGACATCAGGAGCCCATCGTGGGCTAGGGAAGAT  62
               ||||||| |||||||||||||||||||||||||||||||||||||||||||||||||||| 
Sbjct  174548  GTGGCCCTGTCCCAGAGCACCACCCGAGACATCAGGAGCCCATCGTGGGCTAGGGAAGAC  174607

Query  63      CCTCCGGGACCTAACGGCCCAGGT  86
               ||||||||||||||||||||||||
Sbjct  174608  CCTCCGGGACCTAACGGCCCAGGT  174631
```

```
Query  81      CCAGGTCTTCCACCCTTGGCCACCTCCCCAGGTGATGCCTGAAGCTCAAGGGACTGTGTC  140
               ||||||||||||||||||||||||||||||||||||||||||||||||||||||||||||
Sbjct  175110  CCAGGTCTTCCACCCTTGGCCACCTCCCCAGGTGATGCCTGAAGCTCAAGGGACTGTGTC  175169

Query  141     CACCCTCAGG  150
               ||||||||||
Sbjct  175170  CACCCTCAGG  175179
```

```
Query  147     CAGGCCCTGCCCGTGGCTCTGGATCGGCGGTCCCCATCAGAGGCCTGGGCTGAGTCCTCA  206
               ||||||||||||||||||| ||||||||||||||||||||||||| ||||||||||||||
Sbjct  179840  CAGGCCCTGCCCGTGGCTCCGGATCGGCGGTCCCCATCAGAGGCCCGGGCTGAGTCCTCA  179899

Query  207     GGGGT  211
               || ||
Sbjct  179900  GGTGT  179904
```

```
Query  206     AGGGGTCGACGCAGGTCCTCTGGGCAGCATGGGGTGGTGCCGGGTGGACCCCACGGTGGC  265
               ||||||||||||||||||||||||||||||||||||||||||||||||||||||||||||
Sbjct  182126  AGGGGTCGACGCAGGTCCTCTGGGCAGCATGGGGTGGTGCCGGGTGGACCCCACGGTGGC  182185

Query  266     CTGTGTTCGCCACGTGGCCGTGGGTCAGTTCCTCGCTGGAGGAGACCCTTCCCCAGACAC  325
               |||||||| ||||||||||||||||| |||||||||||||||||||||||||||||||||
Sbjct  182186  CTGTGTTCTCCACGTGGCCGTGGGTCGGTTCCTCGCTGGAGGAGACCCTTCCCCAGACAC  182245

Query  326     CTGCCGGGCACCCAGCCAAACCCTTGAGGCCTGGGAAGGTCTCTGGATGCCACCCTACAC  385
               ||||||||||||||||||| ||||||||||||||||||||||||||||||||||||||||
Sbjct  182246  CTGCCGGGCACCCAGCCAAGCCCTTGAGGCCTGGGAAGGTCTCTGGATGCCACCCTACAC  182305

Query  386     CCCGCCCAGACTCCTCTCCCTGTGGTTCCTGGACAGAACATTCCGTCACCACGAAACGAA  445
               ||||||||||||||||||||||||||||||||||||||||||||||||||||||||| ||
Sbjct  182306  CCCGCCCAGACTCCTCTCCCTGTGGTTCCTGGACAGAACATTCCGTCACCACGAAACAAA  182365

Query  446     GTTTGCCCAGGCCCTGCCTTCACCCTGGGAAAACCCAGCCGGGCATGGGAAGGAGGAATG  505
               ||||||||||||||||||||||||||||||||||||||||||||||||||||||||||||
Sbjct  182366  GTTTGCCCAGGCCCTGCCTTCACCCTGGGAAAACCCAGCCGGGCATGGGAAGGAGGAATG  182425

Query  506     CTGCCCGCCTGCCCGCCCAGACCAGAGTCGCCCCACATGCCAGGCCAGGCCCGCATCCTC  565
               ||||||||||||||||||||||||||||||||||||||||||||||||||||||||||||
Sbjct  182426  CTGCCCGCCTGCCCGCCCAGACCAGAGTCGCCCCACATGCCAGGCCAGGCCCGCATCCTC  182485

Query  566     CTGGGGCACACGGGAGCCCCAGGCTTCCCGGCAGGAAAAGCCCCCCACCGCAGAAGGGCT  625
               ||||||||||||||||||||||||||||||||||||||||||||||||||||||||||||
Sbjct  182486  CTGGGGCACACGGGAGCCCCAGGCTTCCCGGCAGGAAAAGCCCCCCACCGCAGAAGGGCT  182545

Query  626     GGTCTCTGGTGGGGAGTTGCTGCTGGGTGGGGTGA  660
               |||||||||||||||||||||||||||||||||||
Sbjct  182546  GGTCTCTGGTGGGGAGTTGCTGCTGGGTGGGGTGA  182580
```
