## Supplementary Figure 5 for "Proteome-wide serology reveals immune-defined subtypes of gastrointestinal disease in systemic sclerosis"

## IP_4486246

**Type** | AltProt
**Gene** | MIRLET7BHG, ENSG00000197182.15
**Organism** | Homo sapiens
**Genomic Coordinates** | chr22:46080338-46105860
**Amino Acids** | 325 (34.74kDa) aa sequence
**External Sources** | None

**Mass Spectrometry**[i]

| studies | unique peptides | detected coverage |
|---|---|---|
| 2 | 2 | 3.7% |

download list of PSMs

| Study | Peptide | Spectrum |
|---|---|---|
| PXD004424 | SSIFLVLR | 151218_exo3_3.9836.9836.2 |

**PSM score** | 0.0856788
**PEP** | 1.61e-1
**experiment** | 151218_exo3_3

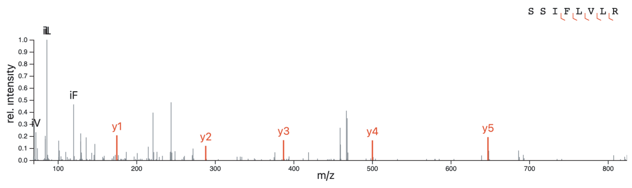


**Mass Spectrometry**[i]

| studies | unique peptides | detected coverage |
|---|---|---|
| 2 | 2 | 3.7% |

download list of PSMs

| Study | Peptide | Spectrum |
|---|---|---|
| PXD005846 | SSIFLVLR | 00864_F1_R1_P0102545B04_TMT10.16407.16407.2 |

**PSM score** | 0.0564345
**PEP** | 8.73e-2
**experiment** | 00864_F1_R1_P0102545B04_TMT10

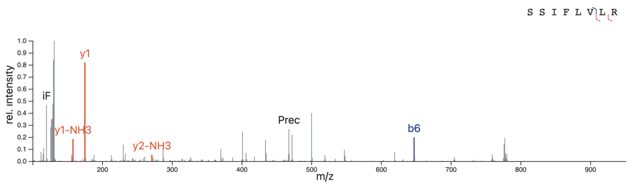


**Mass Spectrometry**[i]

| studies | unique peptides | detected coverage |
|---|---|---|
| 2 | 2 | 3.7% |

download list of PSMs

| Study | Peptide | Spectrum |
|---|---|---|
| BioPlex_2 | TQPGMGRRNAAR | q7712.2494.2494.2 |

**PSM score** | 0.215166
**PEP** | 1.27e-1
**experiment** | q7712

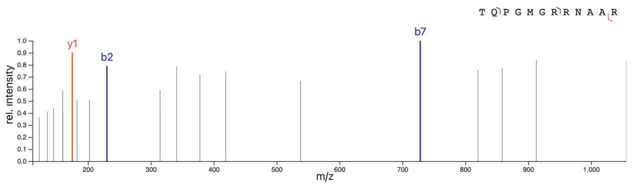


**Mass Spectrometry**[i]

| studies | unique peptides | detected coverage |
|---|---|---|
| 2 | 2 | 3.7% |

download list of PSMs

| Study | Peptide | Spectrum |
|---|---|---|
| PXD004859 | SSIFLVLR | 150828-CTRL2-1-Anus-Remi-F03.18307.18307.2 |

**PSM score** | 0.487842
**PEP** | 6.30e-2
**experiment** | 150828-CTRL2-1-Anus-Remi-F03

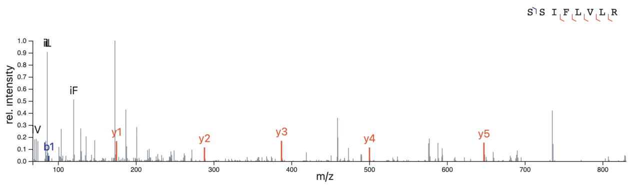


**Mass Spectrometry**[i]

| studies | unique peptides | detected coverage |
|---|---|---|
| 2 | 2 | 3.7% |

download list of PSMs

| Study | Peptide | Spectrum |
|---|---|---|
| TCGA_COCA | AGDKSSIFLVLR | 7843 |

**PSM score** | 0.0144872
**PEP** | 1.22e-1
**experiment** | TCGA-AA-A00R-01A-31_W_VU_20121228_A0218_8E_R_FR08

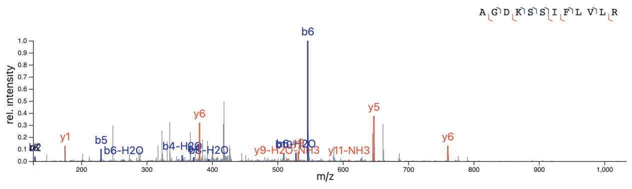


**Mass Spectrometry**[i]

| studies | unique peptides | detected coverage |
|---|---|---|
| 2 | 2 | 3.7% |

download list of PSMs

| Study | Peptide | Spectrum |
|---|---|---|
| PXD004816 | LGPGTSFSLPGFGEDASILR | 160708_46_SC_Human_CFUE_B3_02.69718.69718.3 |

**PSM score** | 0.0051935
**PEP** | 5.06e-1
**experiment** | 160708_46_SC_Human_CFUE_B3_02

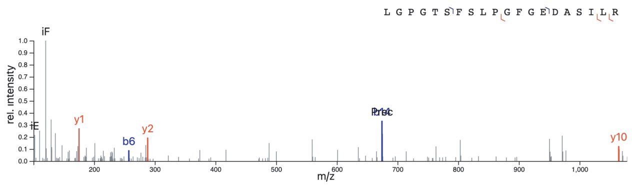


## IP_296890

**Type** | AltProt
**Gene** | LOC124905135, MIRLET7BHG, ENSG00000197182.15
**Organism** | Homo sapiens
**Genomic Coordinates** | chr22:46112549-46112758
**Amino Acids** | 69 (7.74kDa) aa sequence
**External Sources** | None

**Mass Spectrometry**[i]

| studies | unique peptides | detected coverage |
|---|---|---|
| 1 | 1 | 14.5% |

download list of PSMs

| Study | Peptide | Spectrum |
|---|---|---|
| PXD001889 | MSPLPGFR | 20130122_EXQ4_JRW_B71_1_T.20963.20963.2 |

**PSM score** | 0.545972
**PEP** | 4.98e-2
**experiment** | 20130122_EXQ4_JRW_B71_1_T

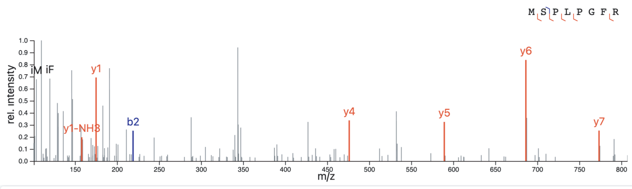


**Mass Spectrometry**[i]

| studies | unique peptides | detected coverage |
|---|---|---|
| 1 | 1 | 14.5% |

download list of PSMs

| Study | Peptide | Spectrum |
|---|---|---|
| PXD001889 | MSPLPGFR | 20130122_EXQ4_JRW_B61_2_T.23104.23104.2 |

**PSM score** | 0.333751
**PEP** | 1.07e-1
**experiment** | 20130122_EXQ4_JRW_B61_2_T

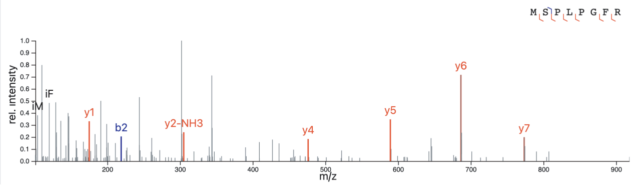


## IP_296881

**Type** | AltProt
**Gene** | LOC124905135, MIRLET7BHG, ENSG00000197182.15
**Organism** | Homo sapiens
**Genomic Coordinates** | chr22:46110885-46111124
**Amino Acids** | 79 (8.95kDa) aa sequence
**External Sources** | None

**Mass Spectrometry**[i]

| studies | unique peptides | detected coverage |
|---|---|---|
| 1 | 1 | 19.0% |

download list of PSMs

| Study | Peptide | Spectrum |
|---|---|---|
| TCGA_OVCA | TWDLLQCPSGGGSRTR | 13045 |

**PSM score** | 0.427036
**PEP** | 3.93e-2
**experiment** | TCGA_24-2295_24-2027_29-1768_117C_W_PNNL_B5S3_f17

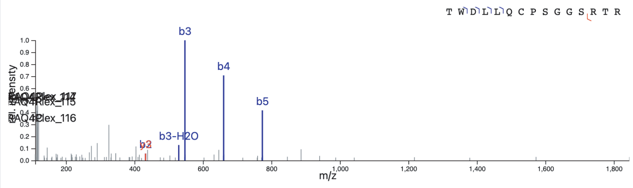
