## Supplementary Figure 7 for "Proteome-wide serology reveals immune-defined subtypes of gastrointestinal disease in systemic sclerosis"

A

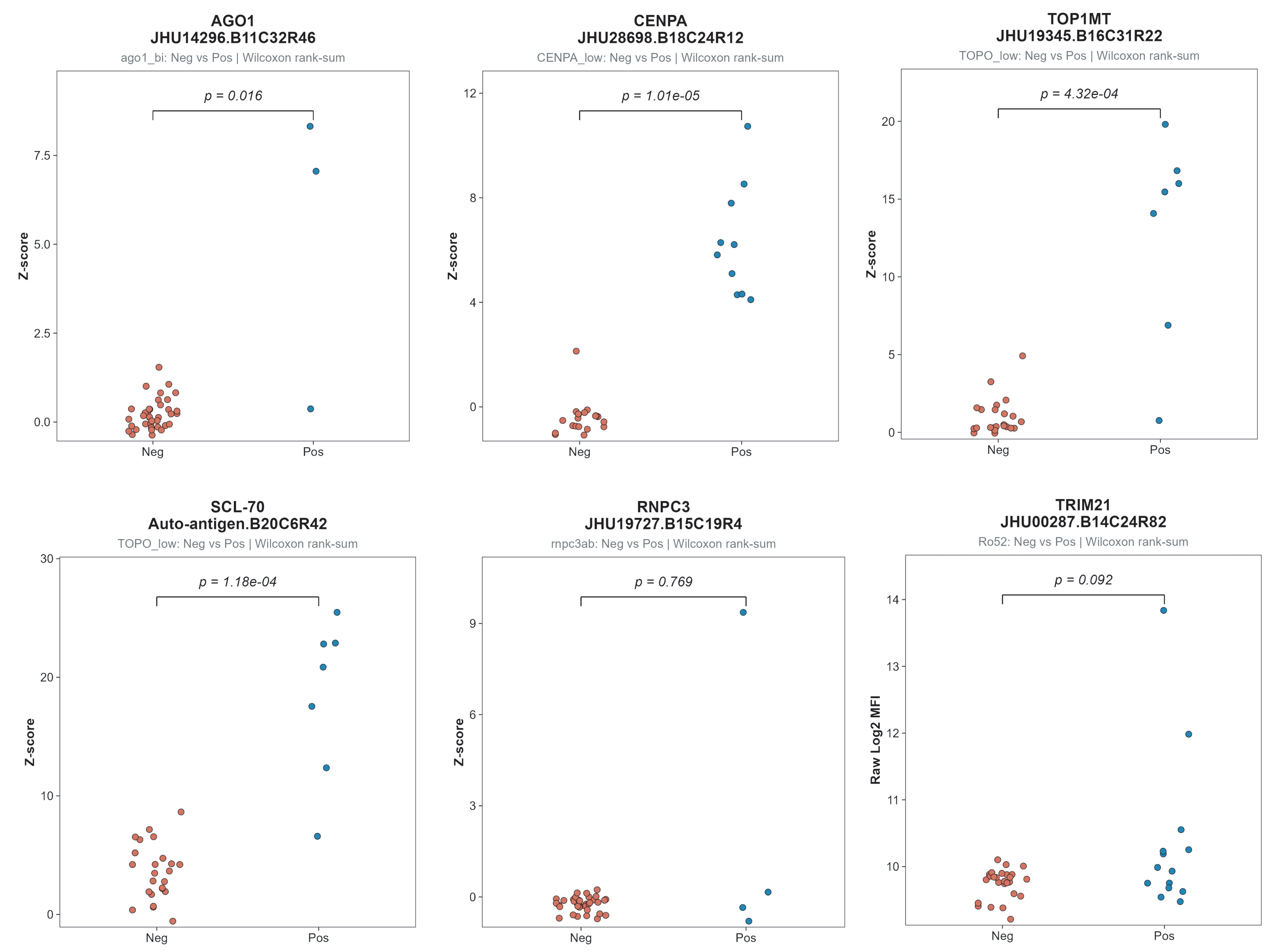


B

| Characteristic | Control (n=20) | Lower_GI (n=17) | Upper_GI (n=23) | Total (n=60) | p-value |
|---|---|---|---|---|---|
| N | 20 | 17 | 23 | 60 | |
| MITO_bi | | | | | |
| Neg | 0 (0.0%) | 15 (88.2%) | 18 (78.3%) | 33 (55.0%) | 0.69 |
| Pos | 0 (0.0%) | 2 (11.8%) | 5 (21.7%) | 7 (11.7%) | |
| Ku | | | | | |
| Neg | 0 (0.0%) | 11 (64.7%) | 19 (82.6%) | 30 (50.0%) | 0.40 |
| Pos | 0 (0.0%) | 1 (5.9%) | 0 (0.0%) | 1 (1.7%) | |
| Ku_low | | | | | |
| Neg | 0 (0.0%) | 11 (64.7%) | 18 (78.3%) | 29 (48.3%) | 1.00 |
| Pos | 0 (0.0%) | 1 (5.9%) | 1 (4.3%) | 2 (3.3%) | |
| PmScl75 | | | | | |
| Neg | 0 (0.0%) | 11 (64.7%) | 19 (82.6%) | 30 (50.0%) | 0.39 |
| Pos | 0 (0.0%) | 1 (5.9%) | 0 (0.0%) | 1 (1.7%) | |
| PmScl75_low | | | | | |
| Neg | 0 (0.0%) | 10 (58.8%) | 18 (78.3%) | 28 (46.7%) | 0.54 |
| Pos | 0 (0.0%) | 2 (11.8%) | 1 (4.3%) | 3 (5.0%) | |
| PmScl100_low | | | | | |
| Neg | 0 (0.0%) | 12 (70.6%) | 18 (78.3%) | 30 (50.0%) | 1.00 |
| Pos | 0 (0.0%) | 0 (0.0%) | 1 (4.3%) | 1 (1.7%) | |
| ThTo | | | | | |
| Neg | 0 (0.0%) | 11 (64.7%) | 19 (82.6%) | 30 (50.0%) | 0.40 |
| Pos | 0 (0.0%) | 1 (5.9%) | 0 (0.0%) | 1 (1.7%) | |
| ThTo_low | | | | | |
| Neg | 0 (0.0%) | 10 (58.8%) | 19 (82.6%) | 29 (48.3%) | 0.16 |
| Pos | 0 (0.0%) | 2 (11.8%) | 0 (0.0%) | 2 (3.3%) | |
| U3RNP_low | | | | | |
| Neg | 0 (0.0%) | 12 (70.6%) | 18 (78.3%) | 30 (50.0%) | 1.00 |
| Pos | 0 (0.0%) | 0 (0.0%) | 1 (4.3%) | 1 (1.7%) | |
| POL155 | | | | | |
| Neg | 0 (0.0%) | 12 (70.6%) | 18 (78.3%) | 30 (50.0%) | 1.00 |
| Pos | 0 (0.0%) | 0 (0.0%) | 1 (4.3%) | 1 (1.7%) | |
| POL155_low | | | | | |
| Neg | 0 (0.0%) | 12 (70.6%) | 18 (78.3%) | 30 (50.0%) | 1.00 |
| Pos | 0 (0.0%) | 0 (0.0%) | 1 (4.3%) | 1 (1.7%) | |
| POL11 | | | | | |
| Neg | 0 (0.0%) | 12 (70.6%) | 18 (78.3%) | 30 (50.0%) | 1.00 |
| Pos | 0 (0.0%) | 0 (0.0%) | 1 (4.3%) | 1 (1.7%) | |
| POL11_low | | | | | |
| Neg | 0 (0.0%) | 12 (70.6%) | 18 (78.3%) | 30 (50.0%) | 1.00 |
| Pos | 0 (0.0%) | 0 (0.0%) | 1 (4.3%) | 1 (1.7%) | |
| CENPB | | | | | |
| Neg | 0 (0.0%) | 8 (47.1%) | 13 (56.5%) | 21 (35.0%) | 1.00 |
| Pos | 0 (0.0%) | 4 (23.5%) | 6 (26.1%) | 10 (16.7%) | |
| CENPB_low | | | | | |
| Neg | 0 (0.0%) | 8 (47.1%) | 13 (56.5%) | 21 (35.0%) | 1.00 |
| Pos | 0 (0.0%) | 4 (23.5%) | 6 (26.1%) | 10 (16.7%) | |
| CENPA | | | | | |
| Neg | 0 (0.0%) | 8 (47.1%) | 13 (56.5%) | 21 (35.0%) | 1.00 |
| Pos | 0 (0.0%) | 4 (23.5%) | 6 (26.1%) | 10 (16.7%) | |
| CENPA_low | | | | | |
| Neg | 0 (0.0%) | 8 (47.1%) | 13 (56.5%) | 21 (35.0%) | 1.00 |
| Pos | 0 (0.0%) | 4 (23.5%) | 6 (26.1%) | 10 (16.7%) | |
| TOPO | | | | | |
| Neg | 0 (0.0%) | 10 (58.8%) | 15 (65.2%) | 25 (41.7%) | 1.00 |
| Pos | 0 (0.0%) | 2 (11.8%) | 4 (17.4%) | 6 (10.0%) | |
| TOPO_low | | | | | |
| Neg | 0 (0.0%) | 9 (52.9%) | 15 (65.2%) | 24 (40.0%) | 1.00 |
| Pos | 0 (0.0%) | 3 (17.6%) | 4 (17.4%) | 7 (11.7%) | |
| rnpc3ab | | | | | |
| Neg | 0 (0.0%) | 17 (100.0%) | 19 (82.6%) | 36 (60.0%) | 0.13 |
| Pos | 0 (0.0%) | 0 (0.0%) | 4 (17.4%) | 4 (6.7%) | |
| Ro52 | | | | | |
| Neg | 0 (0.0%) | 13 (76.5%) | 13 (56.5%) | 26 (43.3%) | 0.32 |
| Pos | 0 (0.0%) | 4 (23.5%) | 10 (43.5%) | 14 (23.3%) | |
| ago1_bi | | | | | |
| Neg | 0 (0.0%) | 15 (88.2%) | 22 (95.7%) | 37 (61.7%) | 0.55 |
| Pos | 0 (0.0%) | 2 (11.8%) | 1 (4.3%) | 3 (5.0%) | |
| ago2_bi | | | | | |
| Neg | 0 (0.0%) | 15 (88.2%) | 22 (95.7%) | 37 (61.7%) | 0.18 |
| Pos | 0 (0.0%) | 2 (11.8%) | 0 (0.0%) | 2 (3.3%) | |
| snrnp35_bi | | | | | |
| Neg | 0 (0.0%) | 17 (100.0%) | 22 (95.7%) | 39 (65.0%) | 1.00 |
| Pos | 0 (0.0%) | 0 (0.0%) | 1 (4.3%) | 1 (1.7%) | |
| U3RNP_bi | | | | | |
| Neg | 0 (0.0%) | 16 (94.1%) | 21 (91.3%) | 37 (61.7%) | 1.00 |
| Pos | 0 (0.0%) | 1 (5.9%) | 2 (8.7%) | 3 (5.0%) | |
| U1RNP_bi | | | | | |
| Neg | 0 (0.0%) | 16 (94.1%) | 18 (78.3%) | 34 (56.7%) | 0.22 |
| Pos | 0 (0.0%) | 1 (5.9%) | 5 (21.7%) | 6 (10.0%) | |
| Platform Coverage | | | | | |
| HuProt_635nm_red | 20 (100.0%) | 17 (100.0%) | 23 (100.0%) | 60 (100.0%) | |
