## Supplementart Data 2 for "Proteome-wide serology reveals immune-defined subtypes of gastrointestinal disease in systemic sclerosis"

We clustered autoantibody profiles and patients based on autoantibodies significantly enriched in lower GI disease compared with controls, which yielded four distinct clusters: $\alpha, \beta, \gamma, \text{and }\delta$.

Cluster $\alpha$ consisted of 18 patients (12 with upper GI disease, 2 with lower GI disease, and 4 controls).

Cluster $\beta$ consisted of 19 patients (3 upper GI, 2 lower GI, and 14 controls).

Cluster $\gamma$ was the smallest, with 6 patients (0 upper GI, 5 lower GI, and 1 control).

Cluster $\delta$ included 17 patients (8 upper GI, 8 lower GI, and 1 control).

Cluster composition revealed clear differences in disease distribution. Cluster $\alpha$ was dominated by upper GI disease patients (66.67%), whereas cluster $\gamma$ was dominated by lower GI disease patients (83.34%). Cluster $\delta$ showed an even representation of upper and lower GI disease (47.05% each), while cluster $\beta$was composed primarily of controls (73.68%).

We also found that autoantibodies against CENPB were highly specific to cluster $\delta.$ A total of 88.23% of cluster $\delta$ patients exhibited high-titer anti-CENPB autoantibodies, whereas no patients in clusters $\alpha$ or $\beta$ had detectable levels of this autoantibody. Only 16.67% (1/6) of cluster $\gamma$ patients were positive, underscoring the strong enrichment of anti-CENPB within cluster $\delta$. Importantly, cluster $\delta$ contains the fewest controls (1/17), which supports the specificity of anti-CENPB for SSc patients. In addition to anti-CENPB, cluster $\delta$ was characterized by elevated frequencies of autoantibodies against CBX1 (76.47%), CBX2 (47.05%), CENPP (76.47%), CENPK (58.82%), DAPK2 (41.17%), EFS (52.94%), CENPA (82.35%), and GGYF1 (52.94%). Anti-CBX1 autoantibodies, for example, were absent in cluster $\gamma$ and detected in only a small proportion of clusters $\alpha$ (11.11%) and $\beta$ (10.52%). Similarly, anti-EFS and anti-GGYF1 autoantibodies appeared in 22.22% and 5.55% of cluster $\alpha$ patients, respectively. Except for these limited occurrences, the autoantibodies listed above were largely specific to cluster $\delta$ and together form a distinct serological signature. Thus, cluster $\delta$, which includes nearly equal proportions of upper and lower GI disease patients, can be defined by a composite serotype characterized by (CBX1, CBX3, CENP-family antigens, DAPK2, EFS, GIGYF1)^+^ autoantibody enrichment.

Cluster $\gamma$ was defined by strong enrichment of autoantibodies against RPP25 (66.67%), RPP30 (50%), POLR3C (50%), LGALS9 (83.33%), LGALS8 (66.67%), and LGALS3 (66.67%). These autoantibodies were either absent or present at substantially lower frequencies in the other clusters: [RPP25 (clusters $\alpha$: 11.11%, $\beta$: 0%, $\delta$: 0%), RPP30 (clusters $\alpha$: 5.55%, $\beta$: 0%, $\delta$: 5.88%), POLR3C (clusters $\alpha$: 5.55%, $\beta$: 0%, $\delta$: 0%), LGALS9 (clusters $\alpha$: 22.22%, $\beta$: 0%, $\delta$: 35.29%), LGALS8 (clusters $\alpha$: 11.11%, $\beta$: 5.26%, $\delta$: 35.29%), and LGALS3 (clusters $\alpha$: 5.55%, $\beta$: 0%, $\delta$: 35.29%). Because LGALS9, LGALS8, and LGALS3 were moderately represented in cluster $\delta$, the distinctive feature of cluster $\gamma$ is the absence of autoantibodies that define cluster $\delta($CBX1, CBX2/CBX3, and centromere proteins) and the marked enrichment of autoantibodies against RPP25, RPP30, POLR3C, and LGALS9. RPP25 (Ribonuclease P/MRP Subunit P25) is a 25 kDa component of the ThTo complex, and RPP30 (Ribonuclease P/MRP Subunit P30), is a 30 kDa component of the same complex. This pattern indicates that anti-ThTo antibodies are a defining serological hallmark of cluster $\gamma$, showing ThTo autoantibodies exclusively in cluster $\gamma$. Thus, the serotype of cluster $\gamma$, which is dominated by patients with lower GI disease, is characterized by (RPP25, RPP30, POLR3C, LGALS9)^+^, in the absence of CBX1, CBX3, CENP family antigens, DAPK2, EFS, and GIGYF1.

Cluster $\alpha$ was defined by an enrichment of autoantibodies against PUF60 (50%) and the astroglial and enteric glial marker GFAP (33.33%). These autoantibodies were not comparably enriched in other clusters [PUF60: $\beta$ 5.26%, $\gamma$ 16.67%, $\delta$ 11.76%; GFAP: $\beta$ 15.78%, $\gamma$: 0%, $\delta$: 17.64%]. Thus, the serotype of cluster $\alpha$, which was dominated by upper GI disease patients, is characterized by (PUF60, GFAP)^+^ in the absence of cluster $\delta$ defining antigens (CBX3, CENPP, CENPA, DAPK2).

In contrast, cluster $\beta$ had the highest proportion of control patients (73.68%) and lacked any distinct autoantibody signature. Although anti-ANXA1 and anti-ARMH4 antibodies were relatively enriched in cluster $\beta$ (36.84% and 73.68%, respectively), both appeared at even higher frequencies in other clusters (e.g., anti-ARMH4 was present in 94.44% of cluster $\alpha$). The predominance of control patients, the absence of serological separation between SSc patients and controls, and the lack of defining autoantibodies indicate that SSc patients in cluster $\beta$ do not share a coherent serotype.

Together, these analyses delineate four serology-based patient clusters, where cluster $\alpha$ (dominated by upper GI disease; defined by PUF60+ and GFAP+), cluster $\gamma$ (dominated by lower GI disease; defined by anti‑Th/To (RPP25/RPP30) and LGALS9+), cluster $\delta$ (balanced upper and lower GI disease; defined by CBX‑family+, CENP‑family+, DAPK2+, EFS+, and GIGYF1+) had cluster-defining serotypes, and cluster $\beta$ (predominantly controls) lacked a definable SSc-associated serotype. These results demonstrate that three of the four clusters (α, γ, δ) possess distinct autoantibody‑based serotypes that align with disease biology, whereas cluster β reflects a heterogeneous group without a serological signature.

Each cluster demonstrated a unique constellation of demographic characteristics, organ involvement, physiologic abnormalities, and autoantibody profiles that we summarize below.

*Cluster* $\alpha$ *[serology of (CBX3, CENPP, CENPA, DAPK2)^-^ AND (PUF60, GFAP)^+^]: Multisystem fibrotic phenotype with severe pulmonary involvement***.** SSc patients in cluster $\alpha$ were middle-aged (mean age 50 ± 14 years) with a median disease duration of 8 years (IQR 5–21), and the most likely to be male (29%) across the clusters. The majority were White (69%) or Black (23%), and diffuse cutaneous disease was present in 50%. Significant Raynaud’s phenomenon occurred in 36% (p = 0.07). This cluster demonstrated the greatest overall disease burden, including the highest proportion of cardiac involvement (25%) and the highest proportion of patients with more severe GI disease by the Medsger score (86%) across clusters. Pulmonary involvement was particularly prominent, as most patients (92%) had pulmonary fibrosis (p = 0.004). Consistent with this, cluster $\alpha$ patients exhibited the lowest forced vital capacity (FVC 67% ± 20; p = 0.004) and the lowest DLCO (45 ± 19; p>0.05). Using previously established serology, cluster $\alpha$ showed the greatest diversity of autoantibody positivity, with frequent anti-Ro52 (43%), anti-U1RNP (29%), anti-RNPC3 (21%), anti-Topoisomerase-1 (20%), and anti-Ago (14%). Despite high physician-assessed GI severity, GI transit testing demonstrated relatively preserved motility, including esophageal emptying at 10 seconds (82% [65–85]) and large-bowel emptying at 72 hours (91% [78–93]). Autonomic symptom burden was moderate, with total COMPASS-31 scores of 33 [21–48].

*Cluster* $\beta$ *[No distinct serology, contains maximum representation of healthy controls]: Vasculopathic and autonomic phenotype.* SSc patients in cluster $\beta$ were younger patients (mean age 47 ± 10 years; p >0.05) with shorter disease duration than clusters $\gamma$ and $\delta$ (8 years [5–31]). Most were white (80%), and none had diffuse cutaneous disease (0%; p = 0.003). In contrast to other clusters, significant Raynaud’s phenomenon was identified in 80% of patients (p = 0.070). Significant GI severity was common (60%), whereas there was no significant cardiopulmonary disease. Pulmonary fibrosis was present in about 50%, but pulmonary function remained relatively preserved on average (FVC 89% ± 16; DLCO 67% ± 16). Right ventricular systolic pressure (RVSP) was the second highest in this group (31 mm Hg [23–32]). Previously known autoantibody profiles included anti-Topoisomerase-1 in 3 patients (60%), anti-RNPC3 in 1 patient (20%), and anti-Ro52 in two (40%). GI motility testing demonstrated slowed esophageal emptying (79% [70–85]) and markedly reduced colonic emptying at 72 hours (44% [0–90]). Concordantly, cluster $\beta$ exhibited the highest total autonomic symptom scores, with the highest COMPASS-31 total scores (50 [22–53]; p>0.05).

*Cluster* $\gamma$ *[serology of (CBX1, CBX3, CENP, DAPK2, EFS, GIGYF1)^-^ AND (RPP25, RPP30, POLR3C, LGALS9)^+^]: Longstanding disease, autoantibody limited, with severe colonic hypomotility (least autonomic symptoms).* Cluster $\gamma$ included the oldest SSc patients (mean age 60 ± 7 years) with the longest disease duration (17 years [10–18]) and was entirely White (100%). Diffuse cutaneous disease was present in 40%, while none had significant Raynaud’s phenomenon or cardiac involvement. 60% had pulmonary fibrosis, though pulmonary function was preserved (FVC 85% ± 8; DLCO 62% ± 21), and RVSP remained low (28 mmHg [25–30]). Previously known autoantibody positivity was infrequent and heterogeneous, including anti-Topoisomerase-1 (20%), anti-Th/To (20%), and anti-Ro52 (20%). GI motility testing revealed the most profound colonic dysfunction across in this cluster, with complete absence of large-bowel emptying at 72 hours (0% [0–0]; p = 0.005). Notably, as expected in a group with severe lower GI disease and relatively normal upper GI disease, this group had the lowest COMPASS-31 scores observed (36 [28–55]).

*Cluster* $\delta$ *[serology of (CBX1, CBX3, CENP, DAPK2, EFS, GIGYF1)^+^]: Anti-centromere–dominant limited cutaneous phenotype with minimal pulmonary fibrosis.* SSc patients in cluster $\delta$ had a mean age of 57 ± 10 years and a disease duration of 12 years (IQR 5–23). This cluster was predominantly white (94%), and all had limited cutaneous SSc (p = 0.003). Raynaud’s phenomenon was present in 33%, and cardiac involvement in 14%. Pulmonary fibrosis was mild relative to other clusters. This group demonstrated the most preserved pulmonary physiology [FVC 91% (15); DLCO 60% (25)] and the lowest prevalence of pulmonary fibrosis (20%; p < 0.001). RVSP was low at 28 mmHg (26–43). In agreement with anti-CENP autoantibodies prevalent in most patients of this cluster, previously known serological profiling also shows that this cluster is dominated by patients with anti-centromere antibodies (91%; p < 0.001). Additional autoantibodies included anti-Ro52 (31%), anti-U1RNP (12%), anti-U3RNP/fibrillarin (12%), and anti-gephyrin (12%), with rare anti-Ku (9%) and anti-Ago (6%) positivity. GI motility was variable but generally preserved, with large-bowel emptying of 71% (19–86). COMPASS-31 scores were the second highest among clusters (40 [24–60]), consistent with mild-to-moderate autonomic symptoms.
