## Supplementart Data 1 for "Proteome-wide serology reveals immune-defined subtypes of gastrointestinal disease in systemic sclerosis"

Unsupervised clustering of autoantibody profiles from all patients, based on autoantibodies enriched in upper GI disease patients compared to control patients was performed. Three broad clusters emerged: clusters A, B, and C.

Cluster A was the smallest, containing 6 patients (4 with lower GI disease and 2 with upper GI disease patients, and no controls).

Cluster C included 17 patients (12 upper GI, 3 lower GI, and 2 controls).

Cluster B was the largest, comprising 34 patients (9 upper GI, 8 lower GI, and 17 controls).

Cluster A was characterized by uniform, high-titer enrichment of anti-CENPB autoantibodies (100%), whereas only 1 patient in cluster B (2.94%), and 9/17 patients in cluster C (52%) were positive for this autoantibody. A similar pattern was observed for anti-CENPA: these autoantibodies were present in all cluster A patients 100%, absent in cluster B (0%), and detected in 47% of cluster C. Autoantibodies against EFS also showed strong enrichment in cluster A (100%), with much lower representation in clusters B (5.88%) and C (29.41%). Thus, CENPA, CENPB, and EFS collectively define the serological signature of cluster A; however, their presence in a substantial proportion of cluster C indicates that these autoantibodies are not exclusive to this group. In contrast, a distinct set of antibodies, including those targeting TEX264, PAG1, SMAD4, NRG3, MUCL1, ST3GAL3, TF, SPAAR, FUT7, SIGLEC8, FUT10, GALNTL6, PXYLP1, PVR, KLRD1, ARMC6, ARRB1, HRAS, LYSMD4, PIEZO1, GCNT2, FAM20B, PACC1, FAM151A, TMEM30A, GCNT1, P2RX4, SLC3A1, PPAT, CHST2, ENPP3, PRPH2, DNAJB6, and SIGLEC5, were enriched in more than 80% of cluster C patients and were either absent or represented at less than 10% in the clusters A and B. These antigens, therefore, delineate a robust cluster C-specific antibody signature.

By comparison, cluster B, the largest cluster, did not harbor any distinctive autoantibody enrichment patterns. Its mixed composition of controls and patients with SSc with both upper and lower GI disease, coupled with the lack of defining serological features, suggests that cluster B does not meaningfully inform disease classification based on autoantibody profiles.
